# Improved detection of differentially represented DNA barcodes for high-throughput lineage phenomics

**DOI:** 10.1101/735266

**Authors:** Yevhen Akimov, Daria Bulanova, Sanna Timonen, Krister Wennerberg, Tero Aittokallio

## Abstract

Cellular DNA barcoding has become a popular approach to study heterogeneity of cell populations and to identify lineages with differential response to cellular stimuli. However, there is a lack of reliable methods for statistical inference of differentially responding lineages. Here, we used mixtures of DNA-barcoded cell pools to generate a realistic benchmark read count dataset for modelling a range of outcomes of lineage-tracing experiments. By accounting for the statistical properties intrinsic to the DNA barcode read count data, we implemented an improved algorithm that provides a significantly higher accuracy at detecting differentially responding lineages, compared to current RNA-seq data analysis algorithms. Building on the reliable statistical methodology, we illustrate how multidimensional phenotypic profiling (or high-throughput ‘lineage phenomics’) enables one to deconvolute phenotypically distinct cell subpopulations within a cancer cell line. The mixture control dataset and our analysis results provide a systematic foundation for benchmarking and improving algorithms for lineage-tracing experiments.

## Introduction

Cellular DNA barcoding was originally developed to trace clonal growth dynamics *in vivo* or *in vitro*^1,2,3,4,5^. More recently, however, cellular DNA barcoding has been applied as an effective means to detect clone-specific differences in the phenotypes other than growth, including drug response^6–12^, postsurgical recurrence^13^, reprogramming capacity^14,15^, phenotypic plasticity^11,16^, and metastatic potential^6,17^. Generally, cellular DNA barcoding can be widely applied to quantify clone-specific differences in virtually any phenotype for which a phenotype-based cell selection method exists. Moreover, emerging methodologies seek to integrate lineage-tracing with single-cell technologies, such as scRNA-seq^15,18–21^, or even isolate lineages carrying a barcode of interest for in-depth cellular profiling^22–24^. These developments are expected to provide high-resolution insights into the biology of heterogeneous cellular systems. However, to our knowledge, there has been no systematic efforts to benchmark the accuracy of clonal phenotype quantification via DNA barcoding.

In a typical clone tracing experiment (Fig. 1A), cells are infected with a short semi-random DNA sequence - a “barcode” - after which the cells are expanded to achieve a sufficient representation of individual clones. The barcoded cell population is then divided into subsamples, typically “control” and “treatment” pools, where the control pool determines a background barcode representation, whereas the treatment pool(s) are subjected to a phenotype-based selection (e.g. drug treatment, immunophenotyping, or xenografting). Finally, the barcode abundances are estimated within each pool with next-generation sequencing (NGS). In the quantification phase, clone sizes are assumed to be proportional to the barcode abundances, and accordingly, differentially represented barcodes (DRBs) between the treatment pool(s) and control population indicate clone-specific differences in the particular phenotype.

**Fig. 1.**
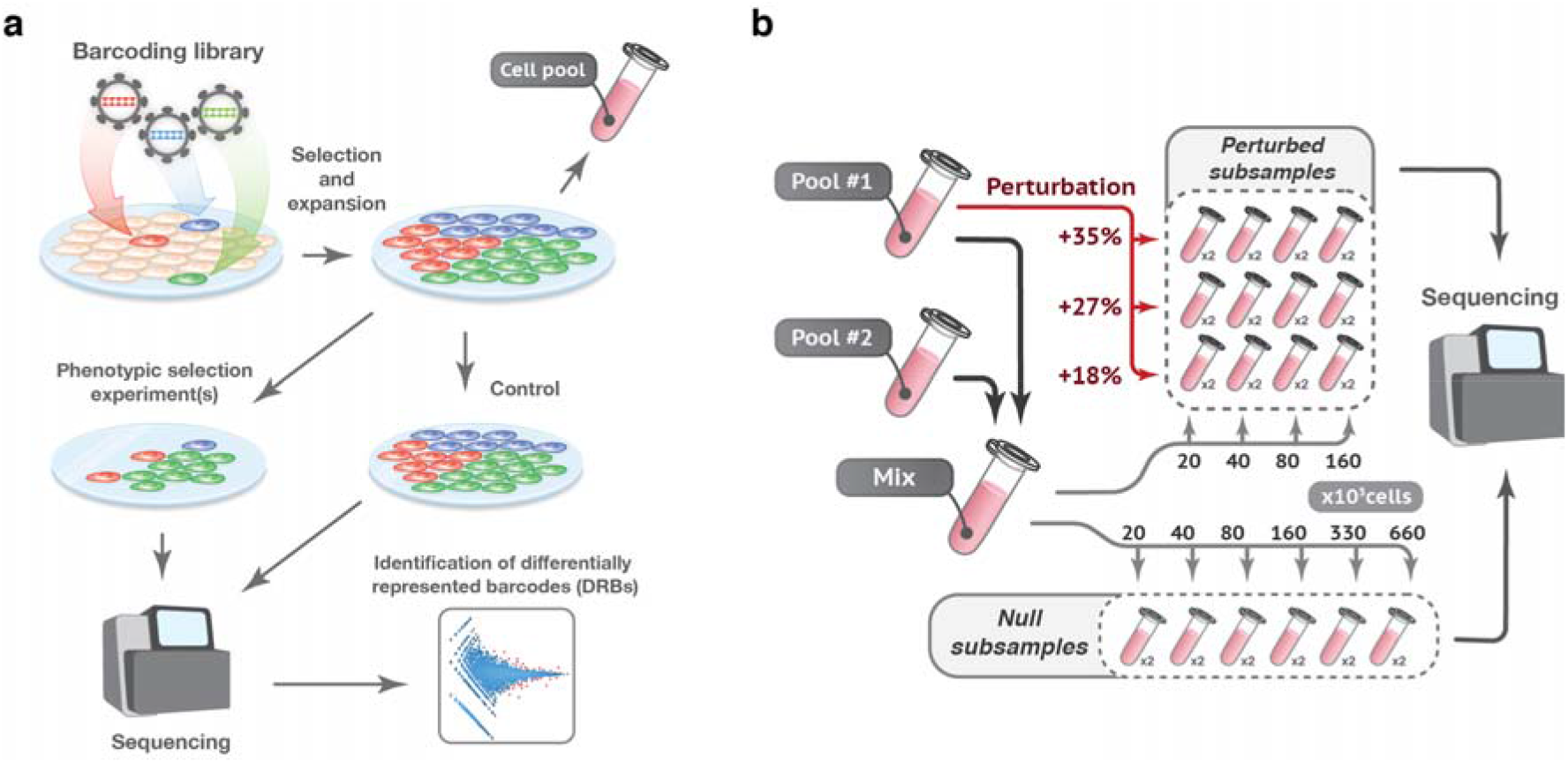
An overview of the experimental setup for the benchmark dataset generation. **a**, A schematic presentation of a typical lineage tracing experiment (see text for description). **b**, To generate the benchmark barcode count datasets, we performed two independent high-complexity DNA-barcoding experiments on Mia-PaCa-II and OVCAR5 cell lines (see Methods for details). In each experiment, cells were collected after selection and expansion step (Fig. 1A) to produce two cell pools (Pool#1 and Pool#2). Cells in each pool were counted and mixed in a 50/50 ratio. The mixture was then subsampled in various extents in two replicas to produce so-called *null subsamples* with different numbers of cells (20×10^3^, 40×10^3^, 80×10^3^, 160×10^3^, 330×10^3^, 660×10^3^) but with the same expected representation of each barcode (i.e., modelling null hypothesis). Perturbed subsamples were generated by taking either 20, 40, 80 or 160 thousand cells from the Pool#1/Pool#2 mixture, and adding an indicated percentages of cells from the Pool#1 (e.g. for sample with 160×10^3^ cells and perturbation degree of 35%, we added 160×10^3^×0.35=56×10^3^ cells from the Pool#1).

The detection of DRBs can statistically be considered as identification of differentially represented sequencing tags from high-throughput count data, and RNA-seq data analysis algorithms have been applied to this task^7^. However, we hypothesized that barcode count data from lineage-tracing experiments may seriously violate the basic assumptions of the RNA-seq analysis algorithms (i.e., that tagwise variance is homogeneous and the read counts follow a negative binomial distribution). We reasoned that the tagwise variance and the underlying distribution of the barcode read counts depends on the sampling size (i.e., the number of sampled cells from the barcoded population). Sampling is an indispensable step in most lineage-tracing experiments. For instance, selection pressure introduces a sampling bottleneck, which reduces the number of cells in the sample proportionally to the degree of the pressure. Such sample size reduction can be extremely high in some applications (e.g., xenografting, high doses of a drug, or cell sorting for rare subpopulations). Therefore, differences in the selection pressure may result in large a variance differences between the samples, leading to biased performance of DRB detection with the RNA-seq analysis algorithms, unless corrected for.

Here, we performed multiple independent clone-tracing experiments on cancer cell lines to generate barcoded cell pools with non-overlapping sets of barcodes. We used these cell pools to generate benchmarking barcode read count datasets for modelling of various outcomes of lineage tracing experiments. By considering the statistical characteristics of the benchmark data and those observed in published studies, we compared the commonly-used RNA-seq analysis algorithms, DESeq^25^, DESeq2^26^, and edgeR^27,28^. Based on the benchmarking results, we developed DEBRA (**DE**Seq-based **B**arcode **R**epresentation **A**nalysis) algorithm for more reliable clone-tracing through improved DRB detection accuracy and a proper control for false discoveries in a wide range of experimental conditions. Finally, we demonstrate how multidimensional phenotypic profiling can be implemented on barcoded cancer cells to identify phenotypically distinct cell subpopulations. These analysis results provide both experimental and statistical insights into high-throughput lineage phenomics and phenotype-based subpopulation inference as an extension of cellular DNA barcoding applications.

## Results

### A gold-standard benchmark dataset for modelling response heterogeneity in lineage tracing experiments

To systematically study the effect of sampling on DNA barcode count data, and the applicability of the RNA-seq data analysis algorithms to the identification of differentially responding lineages, we performed high-complexity cellular DNA barcoding experiments on two cancer cell lines - OVCAR5 and Mia-PaCa-2. The cancer cells were infected with lentiviral DNA barcoding library, carrying ∼5 million unique sequences at very low multiplicity of infection (<0.01 MOI) to label about 5×10^4^ Mia-PaCa-II cells and 10^4^ OVCAR5 cells (see Methods). Each cell line was independently transduced in two replicas, selected with antibiotic and expanded to produce two cell pools with different sets of DNA barcodes (Pool#1 and Pool#2, see Fig. 1). For each cell line, the barcoded cell pools were mixed in a 50/50 ratio and 18 subsamples of different sizes were produced (Fig. 1B, Supplementary Table 1). This experimental setup modelled a situation in which different degrees of selection pressure (e.g. different doses of a drug) are applied to a sample with no lineage-specific differences in the response to the condition (e.g. treatment). We called these samples *null subsamples* because no barcode is expected to be differentially represented and, therefore, an accurate DRB detection algorithm is supposed to accept the null hypothesis for all the barcodes. Such *null subsamples* enabled us to study the effect of sampling size on the statistical characteristics of barcode count data, and to estimate the false discovery rate of DRB detection algorithms.

Furthermore, we generated 24 *perturbed subsamples* by changing the representation of a subset of barcodes in the Pool#1/Pool#2 mixture by adding extra number of cells from the barcoded cell Pool #1 (Fig. 1B, Supplementary Table 1). *Perturbed subsamples* model the selection experiments on a cell population with various degrees of lineage-specific responses to the selection pressure. By sequencing Pool#1 and Pool#2, we determined the ground-truth for differential representation of the barcodes in the perturbed subsamples, which allowed us to assess the accuracy of the DRB detection algorithms.

### Sampling bottleneck affects statistical properties of the DNA-barcode count data and DRB detection accuracy

To investigate the statistical characteristics of the benchmark barcode count data, we first analyzed the mean-variance relationships for each pair of *null subsamples. We* found a marked increase in the variance as the size of the subsample decreases in both OVCAR5 and Mia-PaCa-2 cells (Fig. 2A, B; Supplementary Fig. 1A, B). We observed a similar dependency in the data from a pancreatic cancer patient-derived xenograft (PDX) model published by Seth et al.^7^ (Fig. 2A, B), where the variance of the drug-treated samples is much higher as compared to that of the non-treated controls. The observed difference is likely due to the decrease in the total number of cells (sample size) in response to the drug treatment. We next tested how well the barcode count data follows a negative binomial (NB) distribution using the goodness-of-fit estimation for our OVCAR5 and Mia-PaCa-2 null subsamples and the published pancreatic PDX samples^7^. Notably, the NB model approximated poorly the barcode count data at low count region both in the small-sized OVCAR5 null subsamples and in the PDX drug-treated samples (Fig. 2C; Supplementary Fig. 1C). These properties of the barcode count data violate the basic assumptions made in the RNA-seq data analysis algorithms, which may lead to their sub-optimal performance when applied to DRB detection in lineage tracing experiments.

**Fig. 2.**
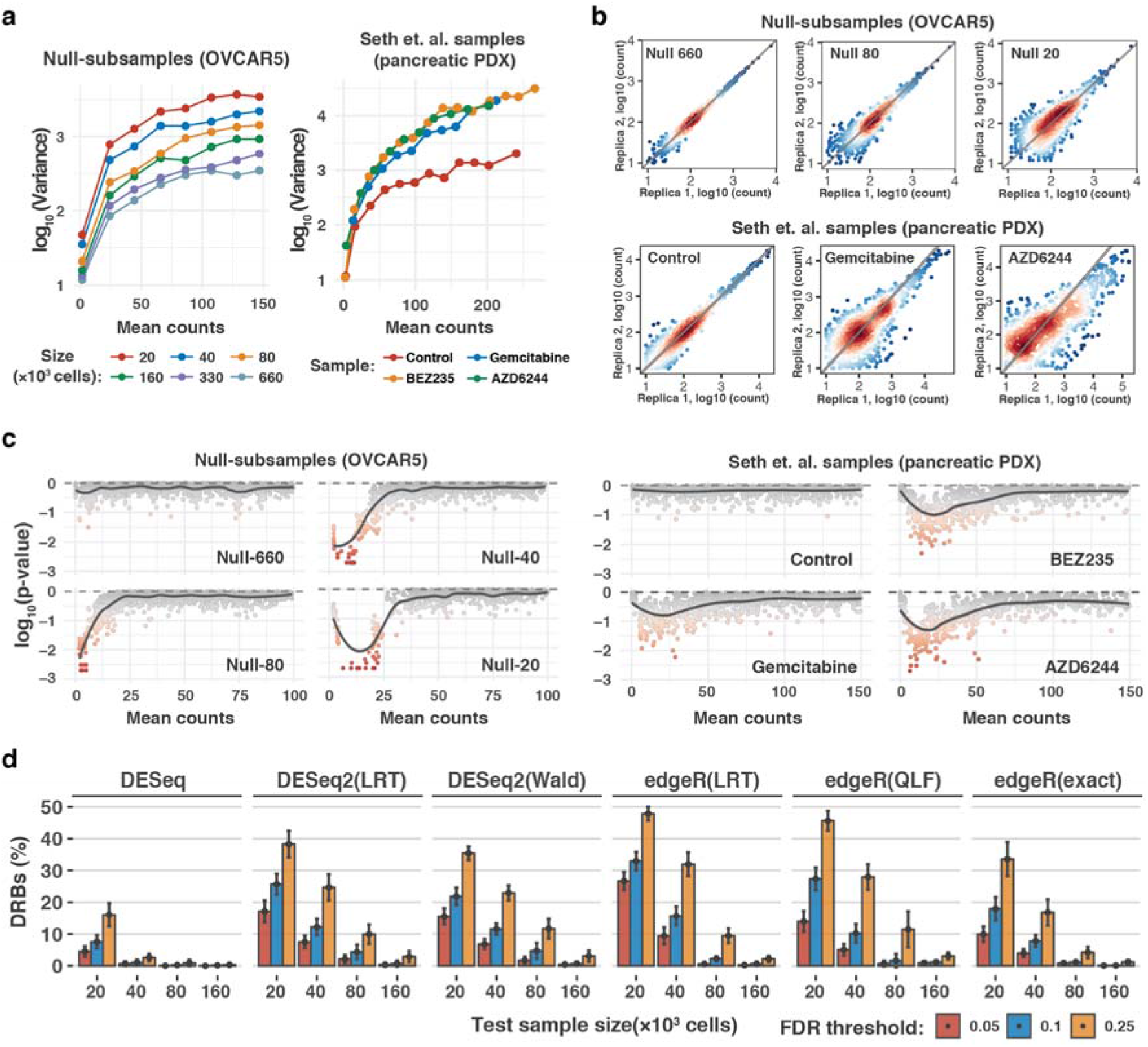
Sampling size affects the statistical properties and accuracy of DRB detection. **a**, Meanvariance plots for the benchmark OVCAR5 null subsamples and pancreatic cancer patient-derived xenograft (PDX) samples^7^. Local variance was calculated by averaging a tagwise variance over the mean counts using a 20 read-count window. **b**, Scatterplots of median-normalized read counts of OVCAR5 null subsamples and pancreatic PDX samples^7^. **c**, Local goodness-of-fit testing for negative binomial distribution where the distribution parameters were estimated using maximum likelihood estimator (MLE). Two-sample Cramer-von-Mises test was used to compare the observed and simulated negative binomial random variables. Statistical significance was determined using Monte-Carlo-bootstrap method, where a small empirical p-value indicates strong deviation from the negative binomial distribution. **d**, The proportion of differentially represented barcodes (DRBs) identified in the OVCAR5 null subsamples with various versions of RNA-seq analysis algorithms. Two replicas of the null subsamples of indicated sizes (x-axis) were tested for DRBs against a control group of 4 null subsamples (two Null-660 samples and two Null-330 samples). The bars represent the average proportion of DRBs identified with the algorithms, calculated over 3-fold bootstrap runs (mean of the 10 resamples with replacement) under the indicted false discovery rates (FDRs). The version with unadjusted p-values is shown in Supplementary Fig. 1D for comparison. LRT, likelihood ratio test; Wald, Wald test; QLF, quasi-likelihood F-test; and exact, implementation of exact test proposed by Robinson and Smyth^29^, as implemented in the original algorithms.

To test the performance of the RNA-seq analysis algorithms for the identification of DRBs, we applied the widely-used algorithms - DESeq, DESeq2 and edgeR - on the OVCAR5 *null subsamples*. An accurate DRB detection method is expected to accept the null hypothesis for all the barcodes (i.e., no barcode should be identified as differentially represented), since the representation of the barcodes are equal across the *null subsamples*. However, all the tested versions of the algorithms identified a significant number of DRBs between the *null subsamples* of different sizes, with percentages of DRBs reaching 50% at smaller sample sizes and higher FDR levels (Fig. 2D). We note that all these detections are false positives, and all the algorithms had much higher type I error rates than those expected based on their empirical p-values (Supplementary Fig. 1D). DESeq performed better than the other algorithms, yet it identified more than 15% false positives at sample size of 20×10^3^ cells at a nominal FDR level of 0.25. Moreover, the performance of DESeq decreased when implemented in other designs (Supplementary Fig. 2A). With all the tested algorithms, the proportion of falsely-detected DRBs increased when *null-subsamples* with larger differences in size and, hence in variance, are compared. These analysis results show that the decrease in sample size due to selection pressure or any other manipulation leading to cell loss may severely compromise the accuracy of DRB detection with the standard RNA-seq analysis algorithms.

### Modified versions of DESeq and DESeq2 algorithms improve the accuracy of DRB detection

We reasoned that the high observed rates of false discoveries is primarily due to the way the RNA-seq analysis algorithms estimate the tagwise dispersions by sharing information across sample groups with unequal variances (e.g. by fitting a negative binomial generalized linear model). To address this issue, we modified the DESeq2 and DESeq algorithms so that the tagwise dispersions are estimated using the replicas of the treatment group only (see Methods). In both of the modified algorithms, we tested two options for tagwise dispersion estimation - “trended” and “shrunken” (see Methods). Due to the observed deviance from the NB model at low-count regions that renders statistical tests based on the NB model inapplicable (Fig. 2C; Supplementary Fig. 1C), we implemented a heuristic algorithm that uses the observed count data to estimate a group-specific read count value (so-called **β** threshold, see Methods), above which the read counts follow the NB model. The estimated **β** threshold was used as a lower bound for the independent filtering step^26,30^ (see Methods). Such an approach effectively eliminates possible false discoveries originating from the read counts that do not follow NB model, while taking advantage of the improved detection power provided by the independent filtering algorithm^26,30^. We implemented the modified DESeq and DESeq2 algorithms into a method dubbed DEBRA (**DE**Seq-based **B**arcode **R**epresentation **A**nalysis), which is available through the CRAN portal (in submission, the script and a workflow example are available through Github https://github.com/YevhenAkimov/DEBRA).

To benchmark the modified algorithms, we first applied DEBRA to the OVCAR5 *null subsamples*. The modified methods correctly accepted the null hypothesis for virtually all the barcodes when the *null subsamples* were tested against each other (Fig. 3A), hence demonstrating a greatly improved control for false discoveries compared to the original algorithms. When the trended dispersion estimates were used, the proportion of identified DRBs were within the range of 0 - 1.5×10^−3^, while the shrunken estimates led to somewhat increased false positive DRB rate of up to 4×10^−3^.

**Fig. 3.**
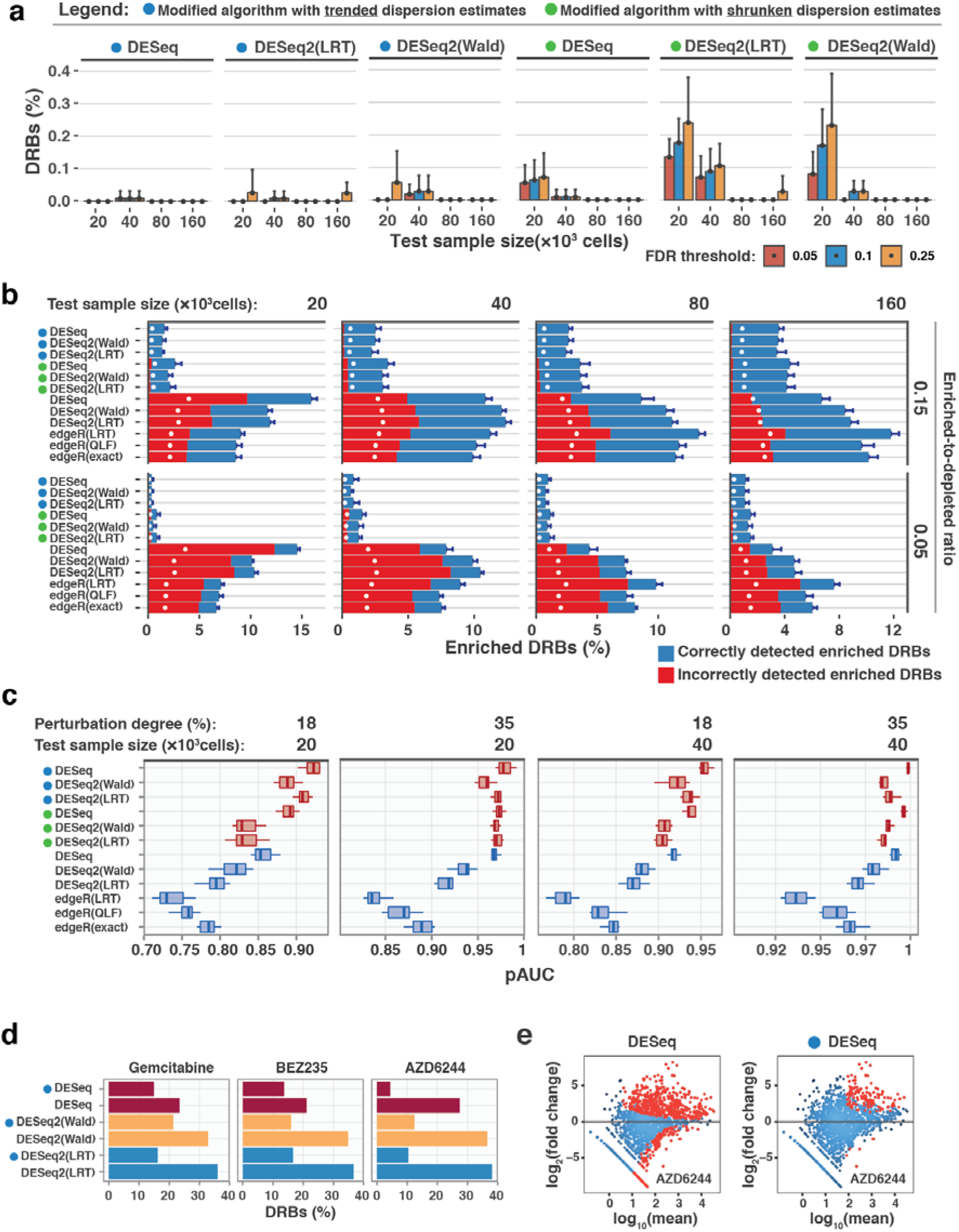
Comparison of the algorithms’ performance. Circles left to the algorithms’ names indicate the modified algorithms. **a**, The percentage of DRBs identified by the modified algorithms in the OVCAR5 null subsamples using the same design as in Fig. 2D. **b**, The performance of the original and modified algorithms for detection enriched barcodes in the perturbed subsamples. Two replicas of the sample with perturbation degree of 35%, indicated size (top) and enriched-to-depleted ratio (right) were tested against four null subsamples (two Null-660 samples and two Null-330 samples). The bars represent the percentage of the barcodes detected as enriched DRBs (fold change > 0; FDR < 0.25) by the indicated algorithm, with correctly detected barcodes marked in blue and incorrectly detected barcodes in red (see Supplementary Fig. 4 for other perturbation degrees and enriched-to-depleted ratios). White circles mark the percentage of barcodes corresponding to the nominal FDR level. **c**, The standardized partial area under the precision-recall curve (pAUC) calculated within intervals of [0,1] and [0,0.25] for precision and recall metrics, respectively. The panel shows the pAUC for perturbed subsamples of indicted size and perturbation degree with enriched-to-depleted barcodes ratio of 0.5 (see Supplementary Fig.s 5 and 6 for pAUCs and precision-recall curves for other sample sizes, perturbation degrees and enriched-to-depleted barcodes ratios). For calculating the precision and recall metrics, we ranked the barcodes with unadjusted p-values as classification scores, where the positive class was defined as correctly detected barcodes (correctly assigned to either enriched or depleted group; see Methods for details). **d**, The proportion of significant DRBs with FDR < 0.1 in the Seth et al. pancreatic PDX dataset^7^, as identified with the modified and original algorithms. **e**, Log fold change vs log mean plots for the AZD6244 drug-treated samples tested against untreated control^7^ with the original DESeq and modified (trended) DESeq algorithms. Red dots mark barcodes with FDR < 0.1.

To test the accuracy of the modified algorithms at detecting DRBs, we used the *perturbed subsamples* to model experimental outcomes with varying proportions of enriched and depleted lineages. The ground-truth for the differential barcode representation in the perturbed subsamples was determined by assigning each barcode to the enriched or depleted group according to its presence either in Pool#1 or Pool#2, as defined by sequencing of the cell pools. The ground-truth information was used to generate experimental results (read count tables), with varying enriched-to-depleted barcode ratios (0.05, 0.15 and 0.5; 10 replicas for each size and perturbation degree; see Supplementary Fig. 3 for details). We tested each modelled experimental outcome for DRBs using the *null subsamples* as a control. The original algorithms showed again relatively high rates of false positives in the low-sized samples with enriched-to-depleted barcodes ratios of 0.05 and 0.15 (Fig. 3b). Notably, the rates of false positives were higher than expected by the nominal FDR levels (Fig 3b, white circles), except for the samples with enriched-to-depleted ratio of 0.5, where the percentage of false positives dropped below the nominal FDR threshold with the original algorithms (Supplementary Fig. 4). However, in the low-sized samples, the rate of false positives detected by the original algorithms became very close to the results obtained when p-values were randomly permuted over the barcodes (Supplementary Fig. 4). This suggests that the empirical significance testing of the original algorithms cannot properly control for the false positives when samples with high difference in variance are being compared. In contrast, the false positive rates of the modified algorithms with trended dispersion estimates never exceeded the nominal FDR threshold, demonstrating an improved control for false discoveries when detecting DRBs in all the tested conditions (Fig. 3B; Supplementary Fig. 4).

To systematically test their accuracy for DRB scoring, we further compared the performance of the algorithms using partial area under precision-recall curve (pAUC) as a summary performance metric (see Methods). In this analysis, the barcodes were ranked by the unadjusted p-values from the algorithms, with low p-values indicating high statistical confidence that the barcode was either enriched or depleted. We found that the modified algorithms with trended dispersion estimates provided better barcode scoring in virtually all the tested scenarios, further supporting its improved performance (Fig. 3C). Among all the tested versions, the modified DESeq with trended dispersion estimates showed the most robust scoring across all the tested conditions (Supplementary Figs. 5 and 6). When applied to the pancreatic PDX data^7^, the modified algorithms with trended dispersion estimates identified again substantially less number of DRBs under the same FDR threshold than the original algorithms (Fig. 3D). Consistently with results from the benchmarking dataset, the difference in the number of detected DRBs between the original and modified methods was larger for the higher-variance sample (AZD6244; Fig. 3E)

### Multidimensional phenotypic profiling of barcoded cells identify distinct cancer cell subpopulations

To further widen the applicability of the lineage tracing technology, we introduce a novel concept of DNA barcoding-based high-throughput lineage phenomics. In this approach, multiple phenotypes are measured for each lineage in a population to produce a multidimensional phenotypic profile for single-lineage analysis. To illustrate this approach, we quantified multiple lineage-specific phenotypes for the barcoded OVCAR5 cell line (Fig. 4A, Supplementary Fig. 7) by applying a number of independent phenotypic assays that introduce differing selection pressure to barcoded OVCAR5 cells, and then analyzed the lineage-specific responses using DEBRA. We first validated the approach with measurements of lineage proliferation rate assessed by two independent readouts, lineage-specific KI67 protein expression^31^ and the number of lineage doublings. As expected, there was a positive correlation between lineage growth rates and lineage KI67 staining (Fig. 4B, Supplementary Fig. 8A), suggesting the feasibility of multidimensional lineage phenotyping via cellular DNA barcoding.

**Fig. 4.**
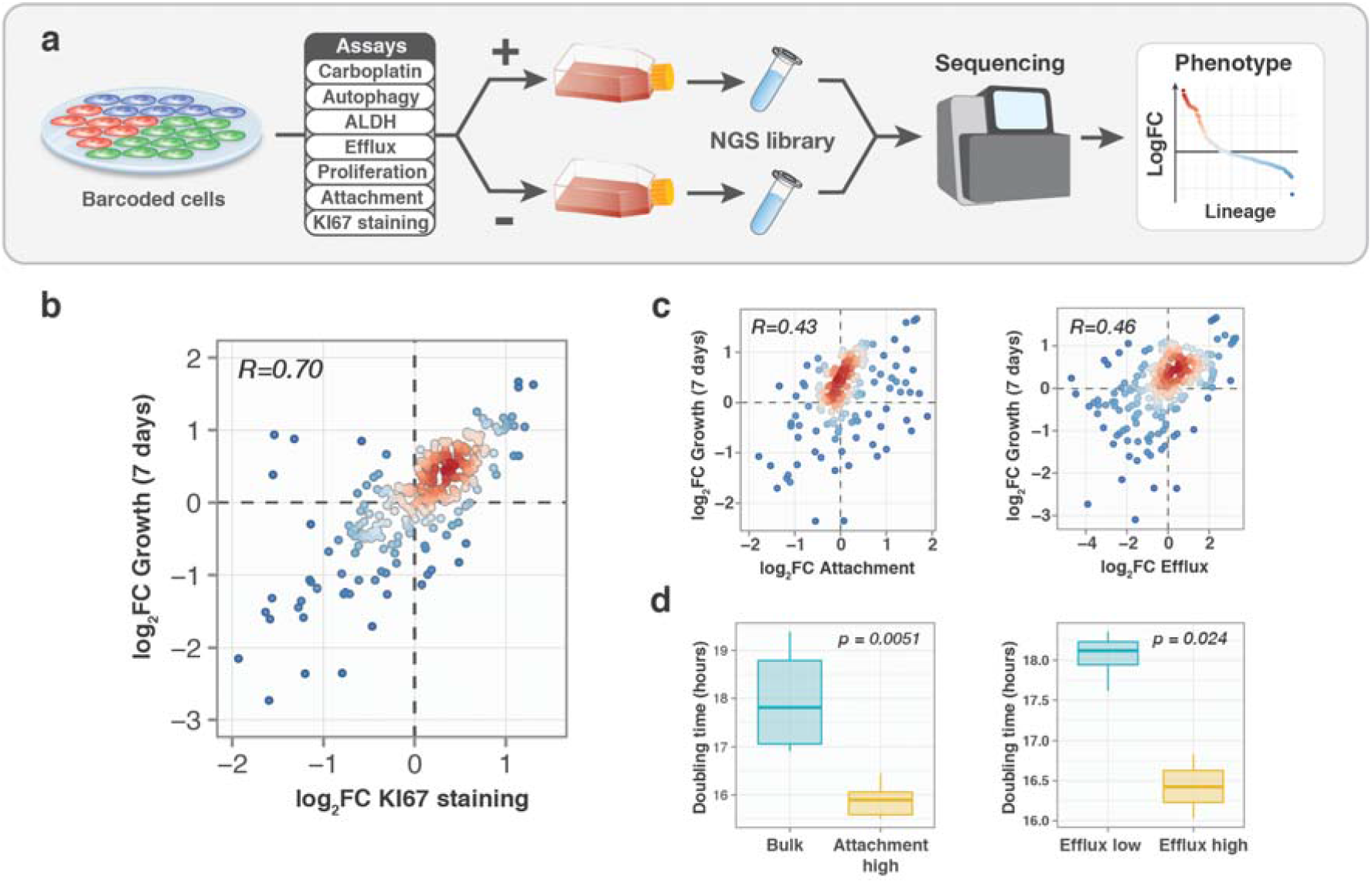
Validation of single-lineage phenotypic profiling approach. **a**, A schematic presentation of the experimental workflow for barcoding-based high throughput multidimensional lineage phenomics approach. Cells were barcoded and expanded to achieve reasonable representation of cells per barcode (e.g. 500-1000). Next, the population was divided into multiple samples and selection pressure was applied to each sample. Cells passing selection conditions were collected and used to prepare a NGS library. In the present study, we measured lineage-specific fold changes in barcode representation in the following assays: carboplatin response (7uM carboplatin for 3 days followed by 4 days regrowth), autophagy measured by autolysosomes load (FACS)^32^, ALDH activity (FACS), activity of efflux pumps (FACS), proliferation (7 days), 12 hours attachment assay in FBS-free media (attached and non-attached cells were collected), and KI67 staining (FACS sorting). **b**, Scatterplot of fold change in the barcode representation after 7 days growth versus fold change in representation between KI67^HIGH^ population and control. Each point represents a lineage with color indicating the local density of points. Displayed are only lineages with counts larger than 70. R, Pearson correlation coefficient. **c**, Scatterplot of barcode fraction fold changes after attachment in FBS-free condition and 7 days growth (left), or upon sorting by efficacy of fluorescent dye efflux and 7 days of growth (right), as described in Methods. **d**, The average doubling time of the phenotypic cell subpopulations separated by their attachment to substrate in FBS-free conditions (left; 6 replicas for each group) or sorted by their efficacy to efflux fluorescent dye (right; 3 replicas for efflux^HIGH^ and 6 replicas for efflux^LOW^). P-values are from Wilcoxon test.

Similarly, we found that the proliferation rate of the OVCAR5 lineages correlated positively with their efflux capacity and the ability of the lineages to attach to the substrate in FBS-free conditions (Fig. 4C). We tested if these phenotypes show the same association on the cell subpopulation level. Since the efflux and attachment assays were non-destructive to cells, we isolated populations of Efflux^HIGH^, Efflux^LOW^ and Attachment^HIGH^ cells, and measured their growth rates. Consistently with the observed correlation on the single-lineage level, the isolated Efflux^HIGH^ and Attachment^HIGH^ subpopulations showed higher proliferation rates when compared to Efflux^LOW^ and bulk OVACR5 cells, respectively (Fig. 4D). These results suggest that a correlation between phenotypes identified at the level of individual lineages predicts phenotype-phenotype relationships at the level of cell subpopulations.

Finally, we used the t-SNE dimensionality reduction algorithm^33^ to deconvolute cell subpopulations based on single-lineage phenotypes measured in OVCAR5 cells. The t-SNE projection enabled us to identify 4 clusters of lineages with distinct phenotypic characteristics (Fig. 5A-C). Interestingly, two of the identified clusters displayed carboplatin resistance phenotype (Fig. 5, clusters 2 and 4). Cells from cluster 2 (∼8% of the population) exhibited an increased efflux capacity which is known to mediate the carboplatin resistance^34,35^. Cells from cluster 4 (about 1.5% of the population) displayed slower proliferation rates, increased ALDH activity, higher autolysosomes load, and resistance to carboplatin (Fig. 5A-C). Such a phenotypic signature is typically attributed to cancer cells with stem-like characteristics^36–40,41,42^. Within the largest cluster of lineages (cluster 1), sensitivity to carboplatin showed a moderate correlation with proliferation rate (Supplementary Fig. 8B). The high-throughput phenotyping of the barcoded lineages suggests that the OVCAR5 cell resistance to carboplatin could emerge through various mechanisms mediated by different cell subpopulations. Together, these data suggests that the high-throughput lineage phenomics approach via DNA barcoding enable one to infer phenotypically distinct cell subpopulations, even within a cell line.

**Fig. 5.**
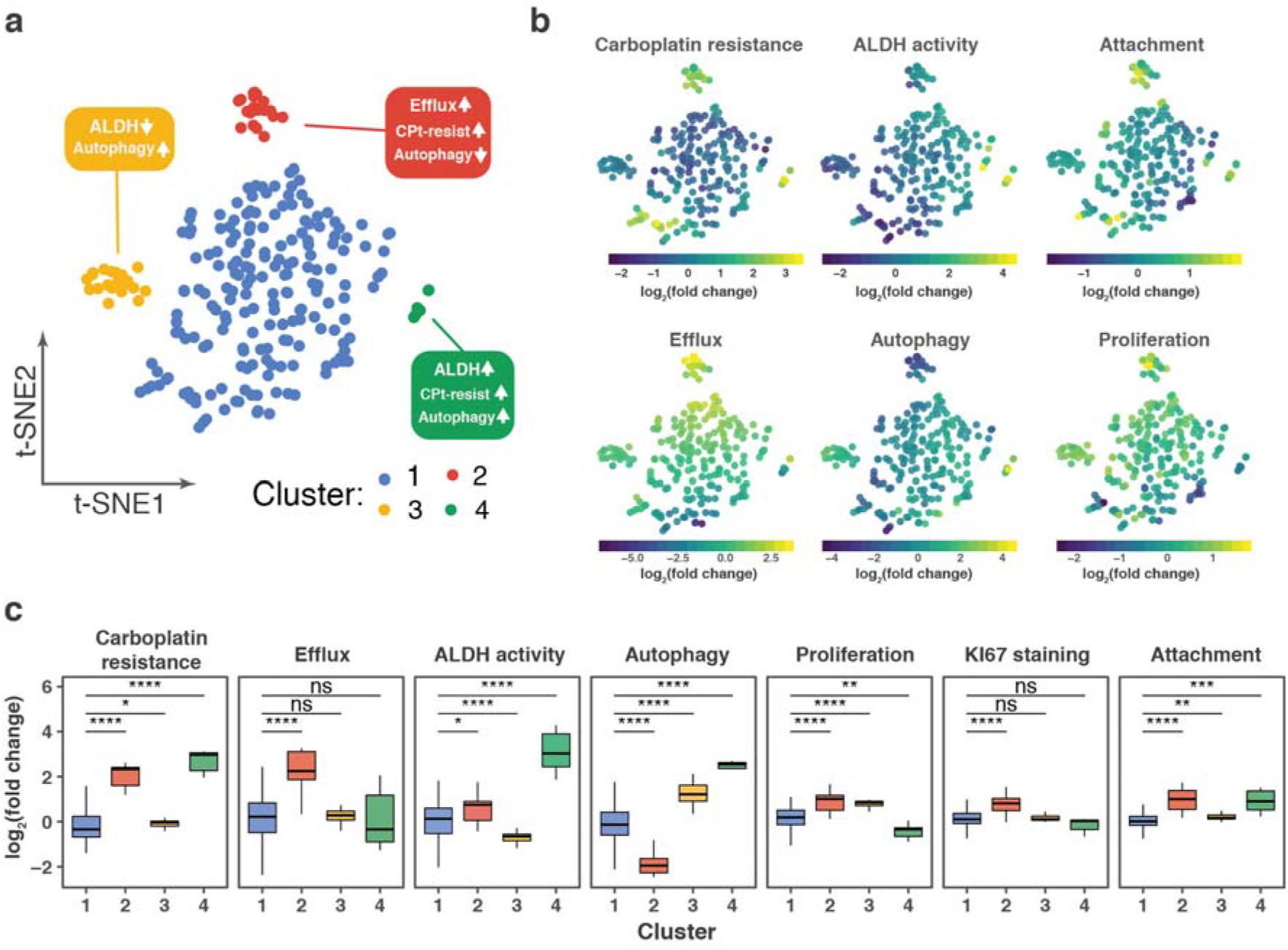
Phenotypic state clustering of single-lineages identifies cancer cell subpopulations. **a**, t-SNE projection of the OVCAR5 single-lineage phenotypic profiles, where each point represents a lineage colored according to the manually gated clusters. The lineages with read counts of more than 75 were used for the statistical analysis. **b**, t-SNE projection of the OVCAR5 single-lineage phenotypic profiles. The lineages are color-coded according to the manifestation of the phenotype, calculated as log2 ratio of barcode fractions between (1) positively and negatively selected populations after ALDH, attachment, efflux capacity or autophagy assays; (2) treated and untreated samples for carboplatin treatment assay, or (3) day 8 and day 1 time points for proliferation assay. **c**, The distribution of log2 fold changes in barcode representations upon selection for the indicated phenotypes. *, p<0.05; ** p<0.01; ***, p<0.001; ****, p<0.0001; ns, non-significant, based on Wilcoxon test.

## Discussion

Lineage tracing via DNA barcoding is a promising method for the studies of intrapopulational heterogeneity of cellular systems. The method is already well-established for tracing growth dynamics of single lineages, but the recent publications suggest a much wider scope of the method’s applicability^6–15^. However, there is a current lack of standardized and reliable frameworks for experimental execution and statistical analysis of the clone-tracing experiments. Towards this end, we implemented a modified algorithm (DEBRA) for reliable detection of DRBs, and demonstrated through systematic benchmarking against the state-of-the-art RNA-seq data analysis algorithms that DEBRA improves the accuracy of detection of differentially responding lineages by accounting for the statistical properties intrinsic to the DNA barcode read count data. The mixture control dataset and our analysis results provide a systematic foundation for benchmarking and improving future algorithms for DNA barcoding data.

Our results showed that samples from lineage tracing experiments exhibit several differences, compared to the RNA sequencing samples, which may compromise the accuracy of DRB identification with the RNA-seq analysis algorithms. In lineage tracing experiments, the number of individual barcodes is close to the total number of cells, which results in around 10^5^ times less of individual sequencing tags as compared to an RNA-seq sample produced from the same number of cells. Hence, we reasoned that any decrease in the cell numbers associated with treatment procedures could impose a sampling error on barcode representation in a manner dependent on the degree of the sample size reduction. Our results support this notion, as we observed strong dependency between sample variance and sample size (Fig. 2). It is also tempting to speculate that the observed deviance from negative binomial model at low counts region is caused by large values of sampling error for barcodes with low copy number. Although we prepared the sequencing libraries right after subsampling, we expect that the variation imposed by the sampling bottleneck is preserved also when the samples are allowed to regrow, something that may have happened in the Seth et al. experiments^7^. We note that increasing the cell expansion times to achieve higher lineage abundances is not a straightforward solution for the sampling error issue. In fact, the expansion time is an indispensable experimental parameter of a lineage tracing experiment, as lineage phenotypes are subject to change as a result of phenotypic plasticity^1,43^. Cellular plasticity may dilute phenotypes determined by non-genetic factors (e.g. epigenetics). Hence, limiting the expansion times is expected to improve quantification of single-lineage phenotypes. Therefore, there is a critical need for accurate detection of DRBs especially in samples with low lineage abundances, using DEBRA or similar algorithmic solutions.

To benchmark the performance of the original and modified algorithms for DRB identification, we simulated lineage tracing experiments with rather challenging scenarios. In the benchmarking mixture cell pool experiments, we used a relatively low number of cells per barcode together with low effects sizes (perturbation degrees of 18, 27 and 35%). These experimental setups are not merely simulated scenarios, in fact many applications of lineage tracing are carried out in the context of a very narrow sample size bottlenecks (e.g. exposure to high doses of drug, xenografting, or cell sorting for rare subpopulations). The DEBRA algorithm was able to both prevent an excess of false discoveries, as assessed with the null and perturbed subsamples, as well as improve the accuracy of DRB classification, compared to the original algorithms, as evaluated with precision-recall analysis (Fig. 3).

We found that DESeq with trended dispersion estimates outperformed the other modified versions, and this is the default option in the DEBRA R-package. The R-package also provides the user with a functionality to choose between two dispersion estimation algorithms - “shrinkage” and “trended” - as the former may be useful in certain experimental setups, e.g., in experiments where the sampling is followed by an extended regrowth. The “trended” method assumes a strict relationship between means and dispersions, whereas the “shrinkage” uses dispersions as estimated by the DESeq2 algorithm. DESeq2 shrinks tagwise dispersion estimates towards dispersion trend using an empirical Bayes approach while allowing for dispersion outliers^26^. In RNA-seq experiments, this helps to deal with genes whose dispersions do not strictly depend on the mean and, therefore, cannot be approximated merely by the dispersion trend. The dispersion outliers are typically attributed to either technical or biological factors. However, it is not clear whether these effects arise also in the lineage tracing experiments. Further studies are needed to better understand the relative benefits of the different dispersion estimation methods in various experimental setups.

Finally, we introduced a DNA barcoding-based lineage phenomics approach, which links multiple phenotypes to lineages in a high-throughput manner (Fig. 4 and Fig. 5). We expect this approach to expedite the inference of cellular subpopulations with distinct phenotypic properties, finding associations between multiple phenotypes and to improve the quantification resolution when analysing intrapopulational phenotypic heterogeneity. The obtained information on the single-lineage phenotypic state could be further integrated with single-cell technologies. For instance, applications of integrated lineage phenomics and single-cell genotyping approaches, such as scRNAseq or scATACseq^44–47^, could promote the discovery of genetic and non-genetic determinants of intrapopulation phenotypic heterogeneity in tumours.

The DEBRA approach could also become useful in the analysis of positive selection CRISPR screens, where the selection pressure is applied to the screening pool (cells expressing Cas9 and sgRNA library), and the representation of sgRNAs in treatment pool is compared to the background distribution. Similar to the DNA barcodes, the sgRNAs may undergo significant representation bottleneck depending on the degree of the selection pressure. Therefore, appropriate control for the variance differences between control and treatment samples, as implemented in DEBRA, may be required for accurate inference of the differentially represented sgRNAs.

## Online Methods

### Generation of the lentiviral plasmid barcode library

Semi-random single-stranded DNA template (Barc.LGMU6.templ; Supplementary Data 3) from Merck (SigmaAldrich) was used in the work. The oligonucleotide was amplified with Barc.LGMU6.aarI.ampl.F and Barc.LGMU6.aarI.ampl.R primers (Supplementary Data 3), using SuperFI DNA polymerase (Thermo, catalog number 12351010) to include cloning overhangs compatible with Golden Gate cloning. Five microliters of the reaction was transferred to a new 50 μl PCR reaction with an excess of Barc.LGMU6.aarI.ampl.F and Barc.LGMU6.aarI.ampl.R primers (Supplementary Data 3). The reaction was run one cycle (2 min at 98°C denaturation, 5 min 72°C annealing/elongation) to produce dsDNA barcodes with no mismatches. The barcode cassette was purified with AMPure XP SPRI beads (Beckman Coulter; catalog number A63880). The barcode cassette was then cloned into previously generated B-GLI-Barcoding plasmid^22^ (see Supplementary Fig. 9 for the plasmid map and Supplementary Data 3 for the DNA sequence; the vector will be deposited to Addgene with publication of the paper^22^) by the Golden Gate assembly method^48^ (see Supplementary Table 2 for reaction composition and cycling conditions). In order to reduce contamination with uncut B-GLI-Barcoding plasmid, an extra 2 μl of the AarI enzyme was added to the reaction after the Golden Gate cycling, followed by incubation at 37°C for 16 h. The cloning reaction was purified with magnetic beads (Beckman Coulter; catalog number A63880) and incubated with Plasmid-Safe™ DNase (Lucigen, catalog number E3101K), according to the manufacturer’s instructions. The reaction was again magnetic beads-purified and transformed into electrocompetent Lucigen Endura™ *E. coli* (Lucigen; catalog number 60242-2) using Bio-Rad MicroPulser Electroporator (catalog number #1652100) with program EC1 following the manufacturer’s instructions. The reaction was plated onto 5 x 15 cm LB-agar plates with 100 μg/ml ampicillin. After incubation for 16 h at 32°C, bacteria were collected and plasmid DNA was extracted with NucleoBond® Xtra Midi kit (MACHEREY-NAGEL; catalog number 740410.50). The efficiency of transformation and approximate number of the unique barcodes in the library was assessed by plating 1/10000 of the reaction onto 15 cm LB-agar plate with 100 μg/ml ampicillin and counting colonies after overnight incubation at 37°C.

### High complexity DNA barcoding experiments

OVCAR5 and Mia-PaCa-2 cells were seeded at a density of 2×10^4^ cells/cm^2^ and 1×10^5^ cells/cm^2^, respectively, both in 6-well plates. Cells were incubated overnight with lentiviral barcoding library carrying ∼5×10^6^ unique barcodes in a presence of 8 mg/ml polybrene. The amount of added virus was selected to achieve a multiplicity of infection (MOI) of ∼0.01. Cells were selected for 7 days in the presence of 150 μg/ml hygromycin. Cells were kept at a density of at least 1×10^4^ cells/cm^2^ to improve viability during selection and expansion.

### NGS library preparation and sequencing

NucleoSpin® Tissue kit (MACHEREY-NAGEL) was used to isolate genomic DNA according to manufacturer’s instructions. Barcodes were amplified from genomic DNA with P5.seq-B-GLI.v1 and P7.seq-B-GLI.v1 primers using OneTaq® DNA Polymerase (NEB; catalog number M0480). Reactions were purified using NucleoSpin® Gel and PCR Clean-up kit (MACHEREY-NAGEL). Then, purified amplicons were amplified with primers, Illumina_indX_F and Illumina_indX_R (where X indicates the index sequence), to add Illumina adapters and indexes for sample multiplexing. This round of PCRs was performed using NEBNext® Ultra™ II Q5® Master Mix (NEB, catalog number M0544). Samples were purified using AMPure XP beads (Beckman Coulter; catalog number A63880). Next generation sequencing library was sequenced with HiSeq 2500 Illumina sequencer using 100 bp paired-end protocol (with 10% PhiX DNA spike-in). To improve cluster calling, we increased sequence diversity by using a 15 bp random sequence stagger in the P5.seq-B-GLI.v1 primer.

### Barcode retrieval from NGS data

We used the previously developed^22^ custom Python script for retrieving original barcode counts from FASTQ files.

### Running DESeq, DESeq2 and edgeR

Dispersion estimation in DESeq^25^ and DESeq2^25,26^ algorithms was implemented using fitType=”local” parameter, as the “parametric” fit option resulted in frequent errors, possibly due to the statistical properties of the barcode count data. Furthermore, we used *method=”per-condition”* setting in DESeq algorithm. The in-built independent filtering option was switched off in DESeq2. The edgeR algorithm was run with its default parameters^28^.

### DEBRA implementation aspects

#### The β threshold

The DEBRA algorithm identifies a threshold **β** - a lower count limit for an independent filtering step above which it is assumed that the read counts follow a negative binomial distribution. This threshold is used for removing results for barcodes with read counts not following negative binomial model and, hence, possibly incorrectly classified as differentially represented. To find a suitable **β** for a given data, the DEBRA algorithm samples read count data using a window of N barcodes ordered by their mean count values (Supplementary Fig. 10). For each sampling step, the algorithm estimates the parameters of the negative binomial (NB) distribution - dispersion (a) and mean (m). DEBRA uses these parameters to generate NB random variables X∼NB(m,a) of the same size as the sampled data to calculate theoretical (expected) and empirical two-sample Kolmogorov-Smirnov (KS) test statistics for each sampling window. The KS empirical test statistic was calculated between the sampled values and X∼NB(m,a) random variables, whilst the theoretical KS statistics is calculated between two X∼NB(m,a) random variables (see Supplementary Fig. 11A for examples). The **β** threshold was estimated by searching for the value of the mean read count at which the overlapping area between the empirical and theoretical density functions of the KS test statistic is close to the maximum overlap for the given data sample. For the estimation, both the theoretical and empirical test statistics are modelled as a Gamma-distributed random variables (see Supplementary Fig. 12) for each window of size N (here, 30 KS test statistics values on the mean ordered data). The overlap area was calculated separately for each window, and then combined from multiple windows by fitting a sigmoid function of mean read counts (see Supplementary Fig. 11B for examples of fitting the null subsamples) with 4 parameters (using drc::drm() function with fct=LL.4() parameter. If the sigmoid curve is ascending and the minimum overlap value is less than 0.25, then **β** threshold is estimated as the mean count at which the sigmoid-fitted overlap takes the value of 0.8 of the maximum (see Supplementary Table 3 for full β threshold estimation rules).

#### Dispersion estimation and inference of differentially represented barcodes

Tagwise dispersions were calculated separately for each condition. We created a DESeqDataSet object, where we pass only the condition-specific columns and calculate the dispersion using DESeq2::estimateDispersions() function using the intercept model (design = ∼1) and fitType=”local” parameter. The trended dispersion estimates were derived from a local dispersion trend function as fitted with DESeq2 (parametrization first proposed in DEXSeq^49^). For calculation, the fitted model were extracted from DESeqDataSet object and used to calculate the tagwise dispersions for the base mean values. The shrunken estimates were extracted from DESeqDataSet object with DESeq2::dispersions() function. The dispersions for barcodes with counts less than **β** in the test samples were set to the maximum value of the calculated tagwise dispersions to reduce false positives from the barcodes not following NB model if the **β** thresholding step is not used. In the next step, the dispersions were passed to the DESeqDataSet (DESeq2) or CountDataSet (DESeq) object, containing full read count dataset (control and condition columns) that are required for inference of DRBs. This object was used to test the barcodes for differential representation with either nbinomWaldTest() or nbinomLRT() tests for DESeq2 implementation or with nbinomTest() for DESeq. Parameter independentFiltering was set to “FALSE” when calling results() function of DESeq2.

#### Independent filtering

We applied the independent filtering procedure^26,30^ as a separate function, which uses DESeq2, DEseq or edgeR result table as an input. The filtering algorithm uses the *genefilter::filtered_p* function to find the number of null hypothesis rejections at a user-specified FDR cutoff (default parameter is set to 0.2) for the quantiles of the filter statistics (mean read counts). The search algorithm identifies the quantile threshold value that maximizes the total number of rejections in the quantile range of [**β**,1], where **β** is the previously estimated threshold for the given data. For the search, the number of rejections is fit as a function of the quantile threshold using a smoothing spline (R function smooth.spline), which enables finding the quantile value that corresponds to the maximum number of rejections. User can also set the **β** threshold value other than the one estimated by the algorithm (see the **β** threshold section).

### Barcode classification and precision-recall curves

A barcode is considered to be differentially represented if the Benjamini and Hochberg procedure-controlled FDR is less than a predefined threshold (here, 0.05, 0.10 and 0.25 were tested). If the count fold change between the treatment and control groups is less than one, then the barcode is considered to be depleted, otherwise it is classified as enriched. Ground truth for the barcode representation in the perturbed subsamples was obtained by sequencing the barcode pools (Pool#1 and Pool#2; see Fig. 1B), which were used to produce the perturbed subsamples. For the ground truth assignments, a barcode is considered enriched if its read Pool#1 to Pool#2 counts ratio is more than 10; if the ratio is less than 0.1, then the barcode is considered depleted. False positive is defined as a barcode identified by the algorithm as enriched DRB, but which is non-enriched according to the ground truth.

Precision-recall curves were constructed using the “precrec” R-package^50^. For calculations, the positive class was defined as a barcode correctly assigned by the algorithm to the group it belongs to (enriched or depleted), while the negative class was defined as a wrongly assigned barcode. We used the unadjusted p-values for the class assignment by the algorithms, i.e., ranking the barcodes against the ground-truth, with low p-values indicating high statistical confidence that the barcode belongs to the positive class (i.e., assigned to either enriched or depleted groups by the algorithm). To calculate the precision-recall metrics for simulated experiments with low enriched-to-depleted barcodes ratios (0.05, 0.15), we used only barcodes with positive fold change values to assess the algorithms’ performance specifically for the enriched barcodes.

### t-SNE algorithm

t-SNE^33^ was run using Rtsne::Rtsne R function^51^ with perplexity parameter of 30 and 1500 iterations. Lineages with read counts > 75 were selected for the analysis.

### OVCAR5 single-lineage phenotypic profiling

Barcoded OVCAR5 cells were grown to reach an average representation of ∼4000 cells per barcode. After that, the pool of cells (5×10^7^) was divided and cryopreserved in 5 batches as a T0 pool. One batch was taken for subsequent phenotypic profiling experiments as outlined in Supplementary Fig. 7.

#### Immunostaining

Cells were trypsinized, washed and resuspended in PBS. Then the cells were fixed and permeabilized with cold 96% ethanol for 30 min on ice, pelleted in a swinging rotor centrifuge at 1000×g for 15 min, rehydrated for 30 min in PBS, washed 2 times in 10 ml of PBS, and blocked in PBS with 0.5% BSA for 1 h at room temperature. The staining was done overnight at 4 C in PBS/BSA. Rabbit anti-Ki67 antibody (ab16667, Abcam) was used at 1.5 μg/ml. Following 3 washes with PBS with 0.5%BSA, the cells were stained with secondary goat-anti-rabbit conjugated with Alexa555 at 1/500 for 30 min at room temperature, washed three times and resuspended in PBS for subsequent sorting.

#### FACS

All the sorting experiments were carried out using SONY SH800Z Sorter at Biomedicum Helsinki FACS Core Facility and the data analysis was performed using Sony Cell Sorter software.

#### ALDH activity assay

The cells in the log phase of growth were trypsinized, resuspended in medium, the concentration of cells was adjusted to 2×10^6^/ml. The ALDH activity was measured using Aldefluor assay (StemCell Technologies, catalog number 01700) according to the manufacturer’s protocol. Cells from the upper and lower quantiles of the Aldefluor fluorescence intensity range were sorted as ALDH^high^ and ALDH^low^ populations, respectively.

#### Efflux assay

Cells were trypsinized, resuspended in medium, and the concentration of cells was adjusted to 2×10^6^/ml. The cells were incubated with CDy1 fluorescent dye diluted 1/1000 (Active Motif, catalog number 895) for 30 min at 37 C in a water bath in the presence or absence of the ABC pumps inhibitors tariquidar (1 μM) and probenecid (50 μM). Then the cells were washed three times in ice-cold PBS and resuspended in medium with or without the drugs. Control cells in medium with efflux inhibitors were left on ice for 2 h, while the test samples were incubated at 37 C for 2 h to allow the efflux of the dye. After 3 washes, the cells were resuspended in PBS, and sorted by the fluorescence intensity in the FL3 (PE-Texas Red) channel. The gating of the efflux-positive cells was set based on the fluorescence intensity of the efflux-inhibited control.

#### Autophagy assay

The autophagy was analyzed by the ratiometric FACS measurement of the amount of Acridine Orange-stained autolysosomes as described previously ^32^. Overnight-starved cells were used as a control for the induction of the autolysosomes formation. 4×10^5^ cells with high autolysosomes load and 10^6^ cells with low autolysosomes load were sorted by FACS for subsequent gDNA extraction.

#### Proliferation assays

For quantification of the cell lineage proliferation rate, the barcoded OVCAR5 cells were propagated in RPMI-1640 medium for 7 passages. Samples for barcode representation analysis were collected at days 0, 5, 8, 11, 14, 18, 25, 29.For the analysis of proliferation rate in validation experiments (Fig. 4D), the cells were plated at 2×10^4^ per well in 12-well plates (Costar), and imaged every 4 h in an IncuCyte HD live cell analysis system (Sartorius) until the cell confluence of all wells reached 100%. The confluence values during the logarithmic growth phase were used to estimate the population doubling time using the formula H/Log_2_(C_F_/C_I_), where H is elapsed time in hours, C_F_ is final confluence, C_I_ is initial confluence.

#### Attachment assay

OVCAR5 cells in the log phase of growth were starved for 16 h in serum-free RPMI supplemented with 2 mM L-glutamine. Upon starvation, the cells were trypsinized, washed in serum-free medium and counted using a Countess II device (Invitrogen). 5 millions of live cells were plated in serum-free medium to 15 cm cell culture dishes and allowed to attach for 12 h. Upon incubation, the non-adherent cells were collected for gDNA extraction by centrifugation at 500×g for 5 min. For the validation experiments, non-adherent cells were collected and replated, and both adherent and non-adherent cells were allowed to recover in serum-supplemented medium for 24 h prior to the evaluation of their proliferation rate.

## Availability of the codes and data

The scripts and a workflow example of the DEBRA implementation are publicly available at Github https://github.com/YevhenAkimov/DEBRA. All the data from the benchmarking and OVCAR5 phenotyping experiments are available in Supplementary Data 1, 2 and 4.

## Funding

This project has received funding from the European Union’s Horizon 2020 research and innovation programme under grant agreement No 667403 for HERCULES. The work was also supported by grants from the Cancer Society of Finland (TA and KW), the Sigrid Jusélius Foundation (TA), Academy of Finland (grants 292611, 279163, 295504, 310507, 313267, 326238 to TA) and Novo Nordisk Foundation (Novo Nordisk Foundation Center for Stem Cell Biology, DanStem; grant NNF17CC0027852 to KW)

## Conflicts of interest

None declared.

## Notes

https://github.com/YevhenAkimov/DEBRA

