## Supplementary Information for "Improved detection of differentially represented DNA barcodes for high-throughput lineage phenomics"

Supplementary Table 1. Details of the benchmark dataset generated using DNA-barcoded cell pools.

| Cell line | Sample name as used in the main text | Number of sampled cells | Number of cells subsampled from 50/50 mix | Number of cells added from pool#1 | Sample name as used in read count tables (Supplementary Data 1 and 2) |
| --- | --- | --- | --- | --- | --- |
| OVCAR5 | Pool#1 | 1E+06 | - | 0 | P1 |
| OVCAR5 | Pool#2 | 1E+06 | - | 0 | P2 |
| OVCAR5 | Null-660 | - | 6,60E+05 | 0 | null_660.1; null_660.2 |
| OVCAR5 | Null-330 | - | 3,30E+05 | 0 | null_330.1; null_330.2 |
| OVCAR5 | Null-160 | - | 1,60E+05 | 0 | null_160.1; null_160.2 |
| OVCAR5 | Null-80 | - | 8,00E+04 | 0 | null_80.1; null_80.2 |
| OVCAR5 | Null-40 | - | 4,00E+04 | 0 | null_40.1; null_40.2 |
| OVCAR5 | Null-20 | - | 2,00E+04 | 0 | null_20.1; null_20.2 |
| OVCAR5 | name not used | - | 1,60E+05 | 5,60E+04 | m_null_160.p35.1; m_null_160.p35.2 |
| OVCAR5 | name not used | - | 1,60E+05 | 4,32E+04 | m_null_160.p27.1; m_null_160.p27.2 |
| OVCAR5 | name not used | - | 1,60E+05 | 2,88E+04 | m_null_160.p18.1; m_null_160.p18.2 |
| OVCAR5 | name not used | - | 8,00E+04 | 2,80E+04 | m_null_80.p35.1; m_null_80.p35.2 |
| OVCAR5 | name not used | - | 8,00E+04 | 2,16E+04 | m_null_80.p27.1; m_null_80.p27.2 |
| OVCAR5 | name not used | - | 8,00E+04 | 1,44E+04 | m_null_80.p18.1; m_null_80.p18.2 |
| OVCAR5 | name not used | - | 4,00E+04 | 1,40E+04 | m_null_40.p35.1; m_null_40.p35.2 |
| OVCAR5 | name not used | - | 4,00E+04 | 1,08E+04 | m_null_40.p27.1; m_null_40.p27.2 |
| OVCAR5 | name not used | - | 4,00E+04 | 7,20E+03 | m_null_40.p18.1; m_null_40.p18.2 |
| OVCAR5 | name not used | - | 2,00E+04 | 7,00E+03 | m_null_20.p35.1; m_null_20.p35.2 |
| OVCAR5 | name not used | - | 2,00E+04 | 5,40E+03 | m_null_20.p27.1; m_null_20.p27.2 |
| OVCAR5 | name not used | - | 2,00E+04 | 3,60E+03 | m_null_20.p18.1; m_null_20.p18.2 |
| Mia-PaCa-2 | Pool#1 (mia) | 1E+06 | - | 0 | Pool1 |
| Mia-PaCa-2 | Pool#2 (mia) | 1E+06 | - | 0 | Pool2 |
| Mia-PaCa-2 | Null-40 (mia) |  | 4,00E+04 | 0 | sampl40_1; sampl40_2; sampl40_3; |
| Mia-PaCa-2 | Null-10 (mia) |  | 2,00E+04 | 0 | sampl10_1; sampl10_2; sampl10_3; |

- stands for “not applicable”

Supplementary Table 2. Golden gate protocol for DNA barcode cloning into B-GLI-Barcoding plasmid

| Golden gate protocol | | |
| --- | --- | --- |
| Rapid Ligation Buffer (Thermo, cat. K1422) | 4 µl | |
| AarI (2U/ul) (Thermo, cat. ER1581) | 1 µl | |
| T4 DNA Ligase (5 U/µL) (Thermo, cat. EL0014) | 0.25 µl | |
| Barcode amplicon | 5 ng | |
| B-GLI-Barcoding | 100 ng | |
| Oligo ( provided with the AarI enzyme) | 0.4 µl | |
| ddH2O | up to 20 µl | |
| Golden gate cycling conditions | | |
| Temperature | Time | Cycles |
| 37C | 5 min | 1 |
| 37C  22C | 5 min  3 min | 50 |
| 37C | 10 min | 1 |
| 65C | 15 min | 1 |
| 4C | hold | 1 |

Supplementary Table 3. **β** threshold estimation rules.

|  | min(overlap)>0.25 | min(overlap)<0.25 |
| --- | --- | --- |
| Descending sigmoid curve | Data follows NB model; beta set to 0 | Visual examination of the results is recommended;  user-specified beta is used |
| Ascending sigmoid curve | Data follows NB model; beta set to 0 | max(overlap)<0.25  Data does not follow NB;  user-specified beta is used  max(overlap)>0.25    beta is estimated as described in methods |

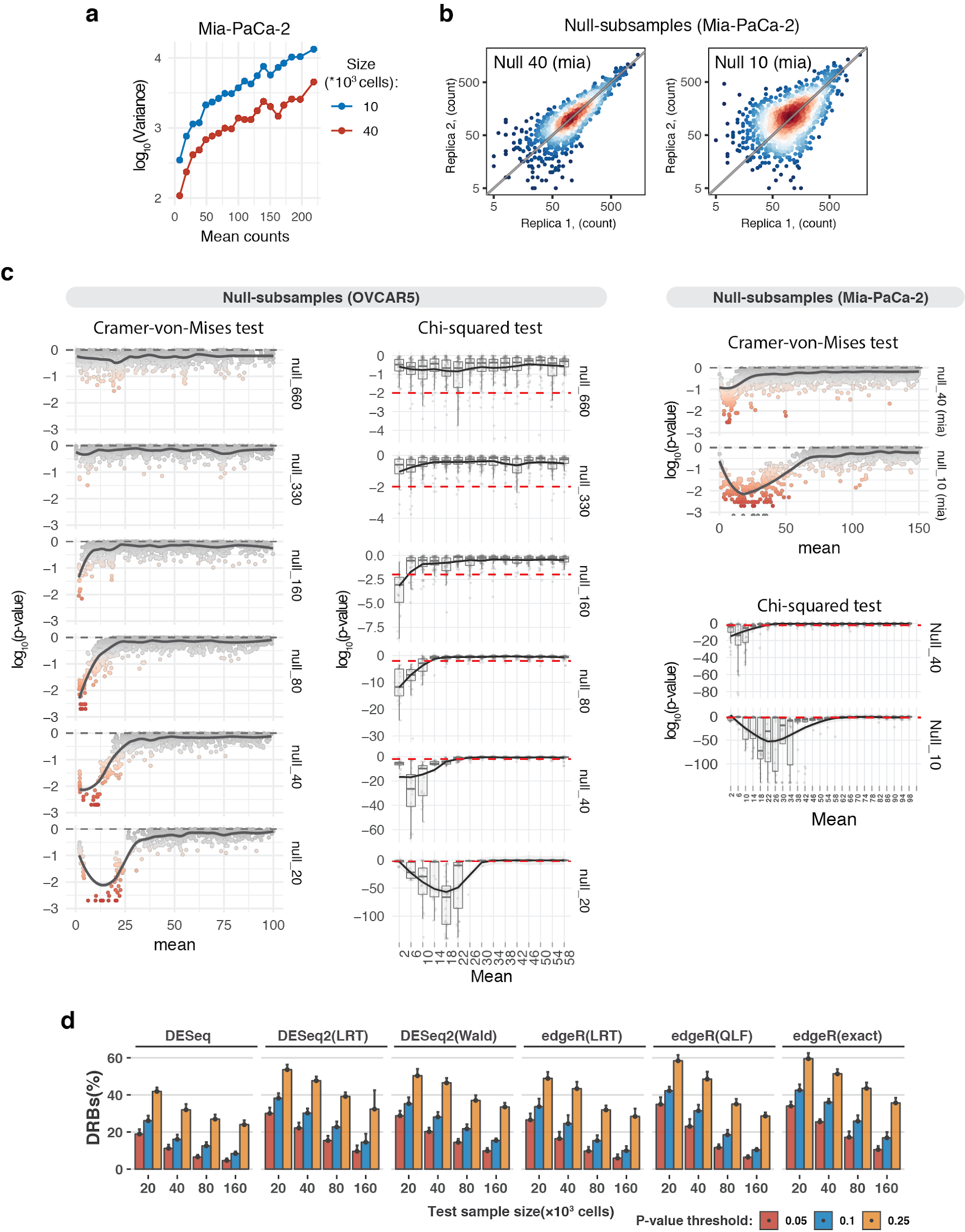

Supplementary Fig. 1. Sampling size affects statistical properties and accuracy of DRB calling.

**a**, Mean-variance plots for Mia-PaCa-2 null subsamples. Barcode read counts were median-normalized. Local variance was calculated by averaging a tagwise variance over the mean counts using a 20 read-count window. **b**, Scatter plots of median-normalized read counts of Mia-PaCa-2 null subsamples. **c**, Local negative binomial goodness-of-fit was estimated using chi-squared test or Cramer-von-Mises test. Dispersion parameter of the negative binomial model was estimated locally over the window of 3 read-counts using maximum likelihood estimator. P-value of the chi-squared test statistics was estimated using fitdistrplus::gofstat() function. P-values of the Cramer-von-Mises test were calculated by Monte-Carlo bootstrap method as implemented in RVAideMemoire::cramer.test. **d**, The proportion of differentially represented barcodes (DRBs) identified in the null subsamples with various RNA-seq analysis algorithms using the same design as in Figure 2D. The bars represent the mean proportion of DRBs calculated over 3-fold bootstrap runs (mean of the 10 resamples with replacement) under indicated unadjusted p-value thresholds.

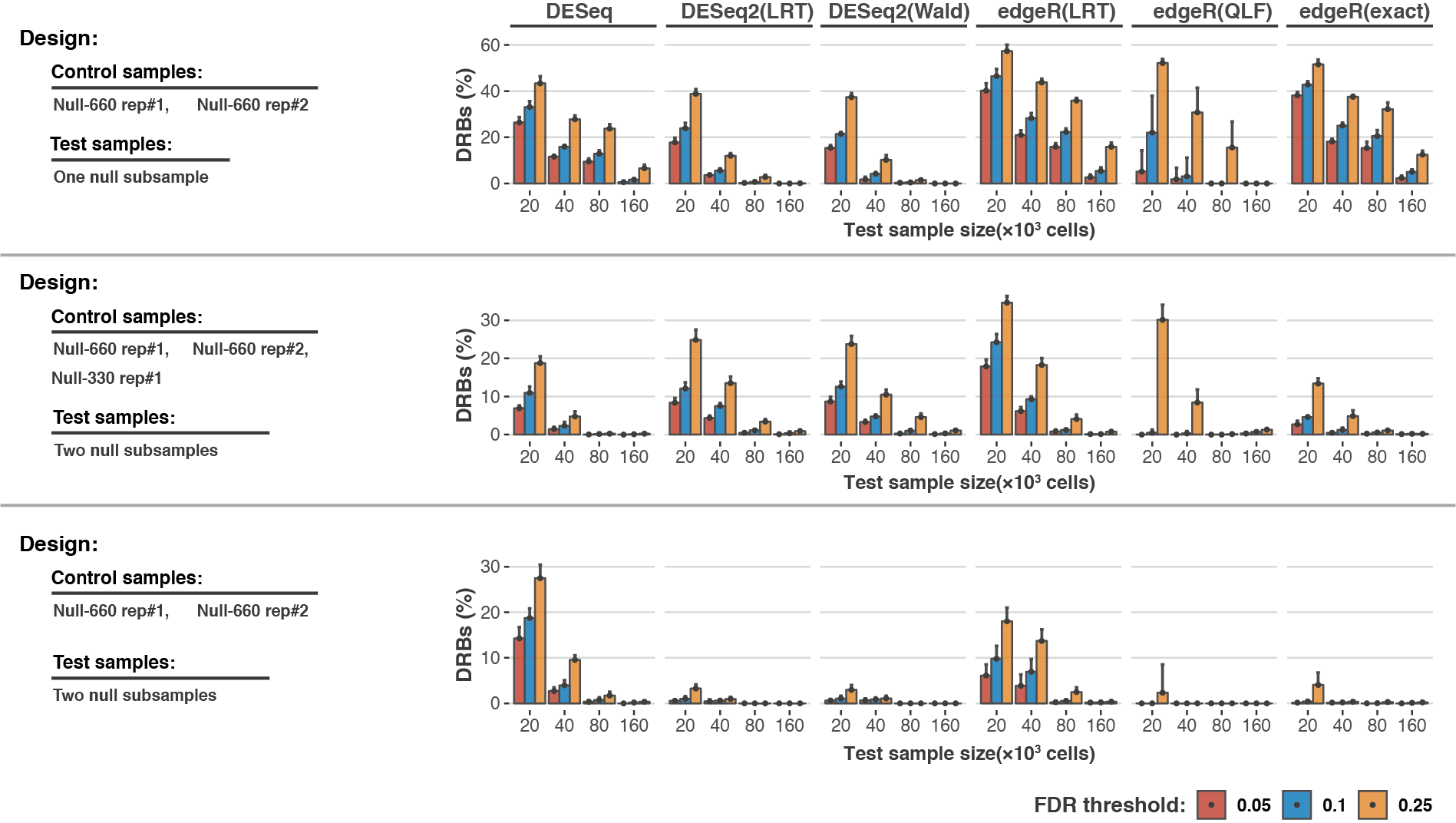

Supplementary Fig. 2. Comparison of the RNA-seq algorithms’ performance on the null subsamples.

Bars represent the proportion of DRBs identified in the OVCAR5 null subsamples with different RNA-seq analysis algorithms. Samples used in control and treatment groups are indicated on the left. The bars represent the average percentage of barcodes called falsely as differentially represented by the indicated algorithms (10 resamplings with replacement), under different nominal false discovery rate (FDRs) thresholds.

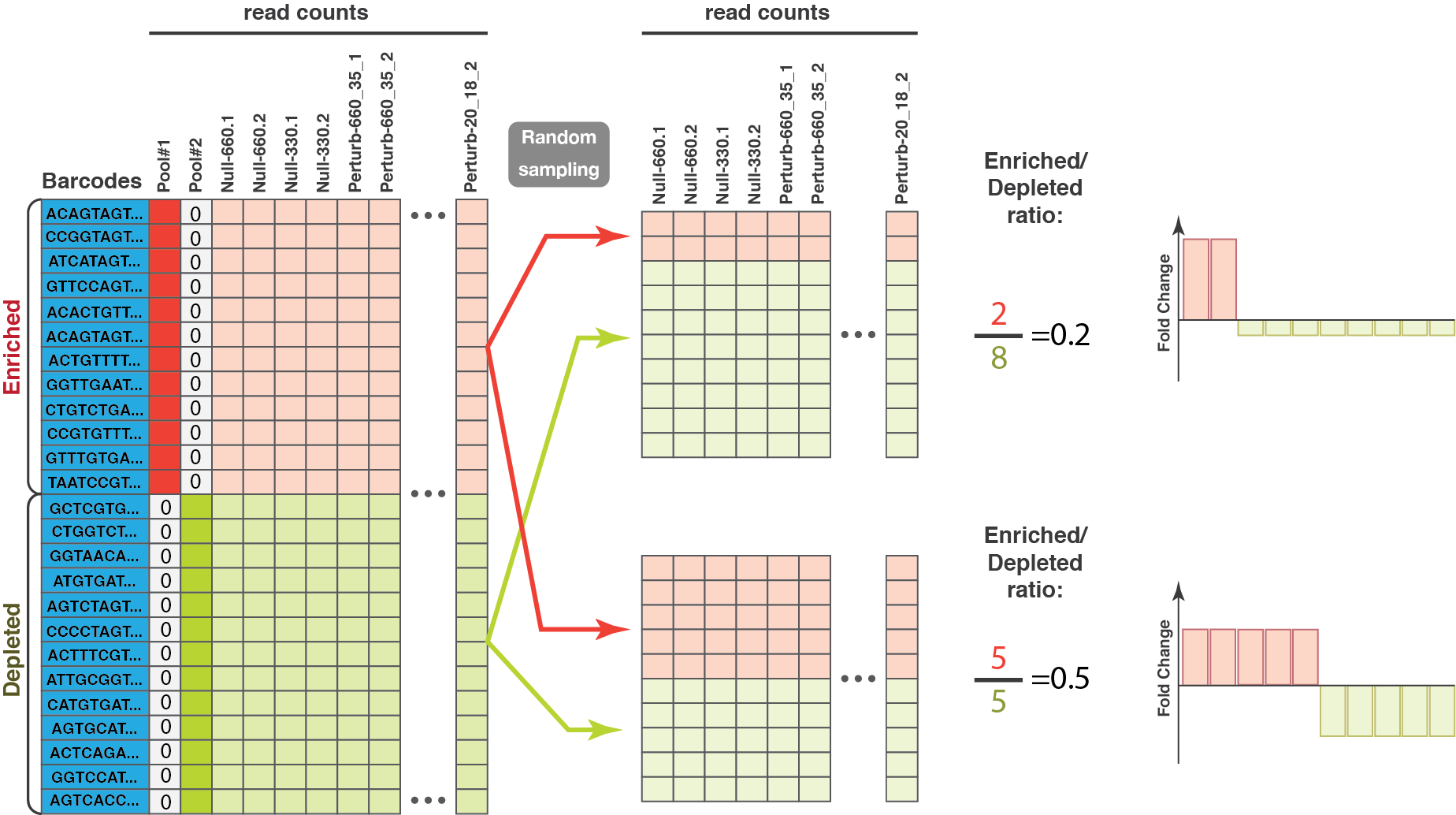

Supplementary Fig. 3. Modelling of barcode experimental results with varying enriched-to-depleted barcode ratios.

The barcodes are assigned into enriched and depleted groups according to the ground-truth (defined by sequencing of the cell pools #1 and #2). Next, the barcodes are sampled without replacement in the desired enriched-to-depleted ratios. Note that after normalization the original perturbation degrees are subject to change, depending on the chosen enriched-to-depleted ratio.

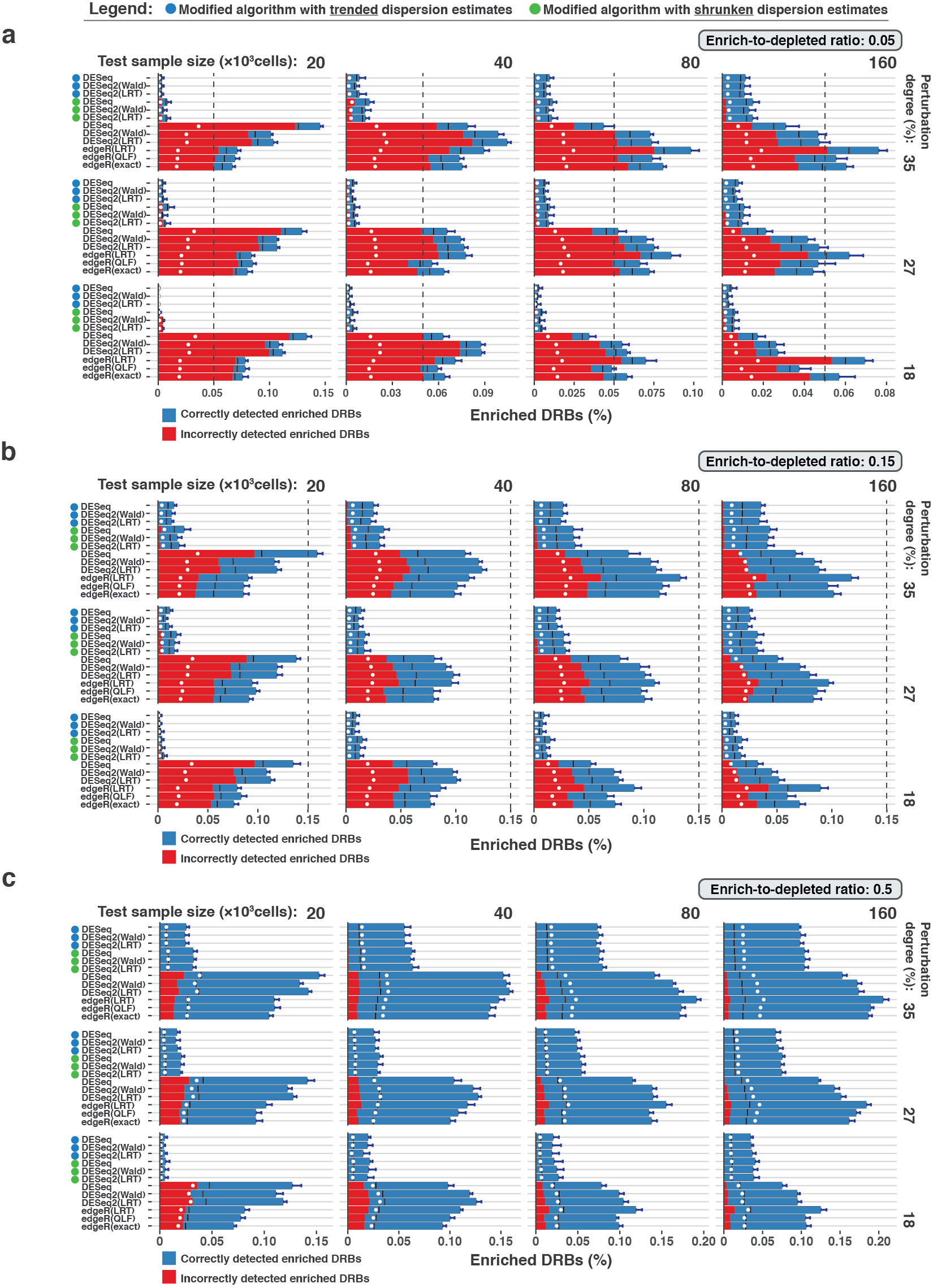

Supplementary Fig. 4. Comparison of the algorithms’ performance on the perturbed subsamples.

Circles left to the algorithms’ names indicate the modified algorithms. Two replicas of the perturbed subsamples of indicated sample size (top), perturbation degree (right) and enriched to depleted barcode ratios of **a**, 0.05, **b**, 0.15 and **c**, 0.5, were tested for DRBs against four control samples (2 Null-660 samples and 2 Null-330 samples). Blue bars represent the proportion of DRBs classified as enriched (fold change > 0) under the FDR threshold of 0.25. Red bars indicate the average fraction of false positives (incorrectly assigned to the enriched group). Black lines indicate the average fraction of false positives observed when p-values were randomly permuted over the barcodes. White points indicate the nominal FDR threshold of 0.25. Dotted vertical lines indicate the total proportion of enriched barcodes.

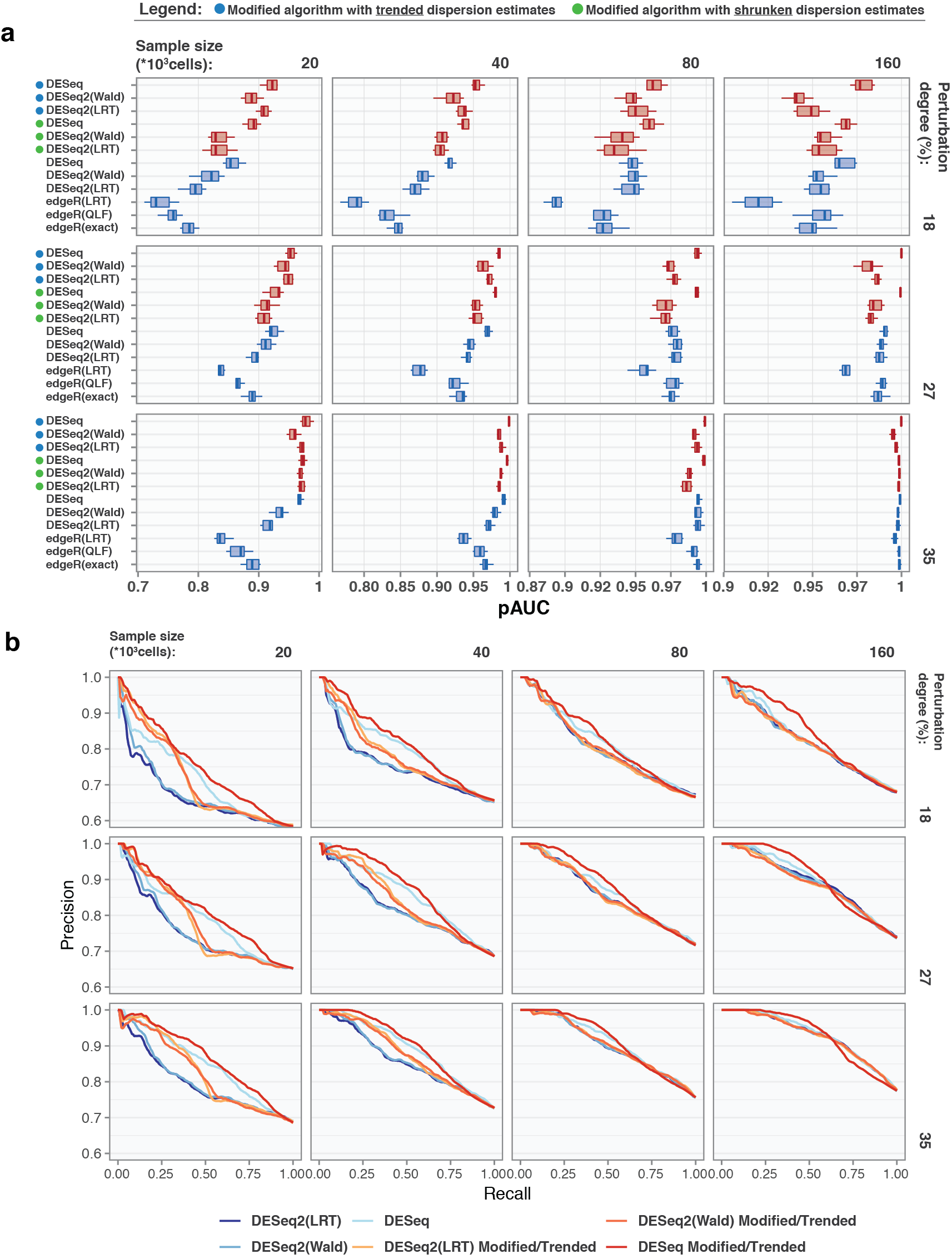

Supplementary Fig. 5. Comparison of the DRB scoring performance using precision-recall analysis.

Circles left to the algorithms’ names indicate the modified algorithms. **a**, Standardized partial area under the precision-recall curve (pAUC) calculated within intervals of [0,1] and [0,0.25] for the precision and recall metrics, respectively. Shown are the pAUCs for perturbed subsamples of indicted size (top) and perturbation degree (right), with enriched to depleted barcodes ratio of 0.5. **b**, Precision-recall curves for the indicated sample sizes (top) and perturbation degrees (right), with enriched to depleted barcodes ratio of 0.5. For clarity, the modified algorithms with shrunken dispersion estimates are not shown.

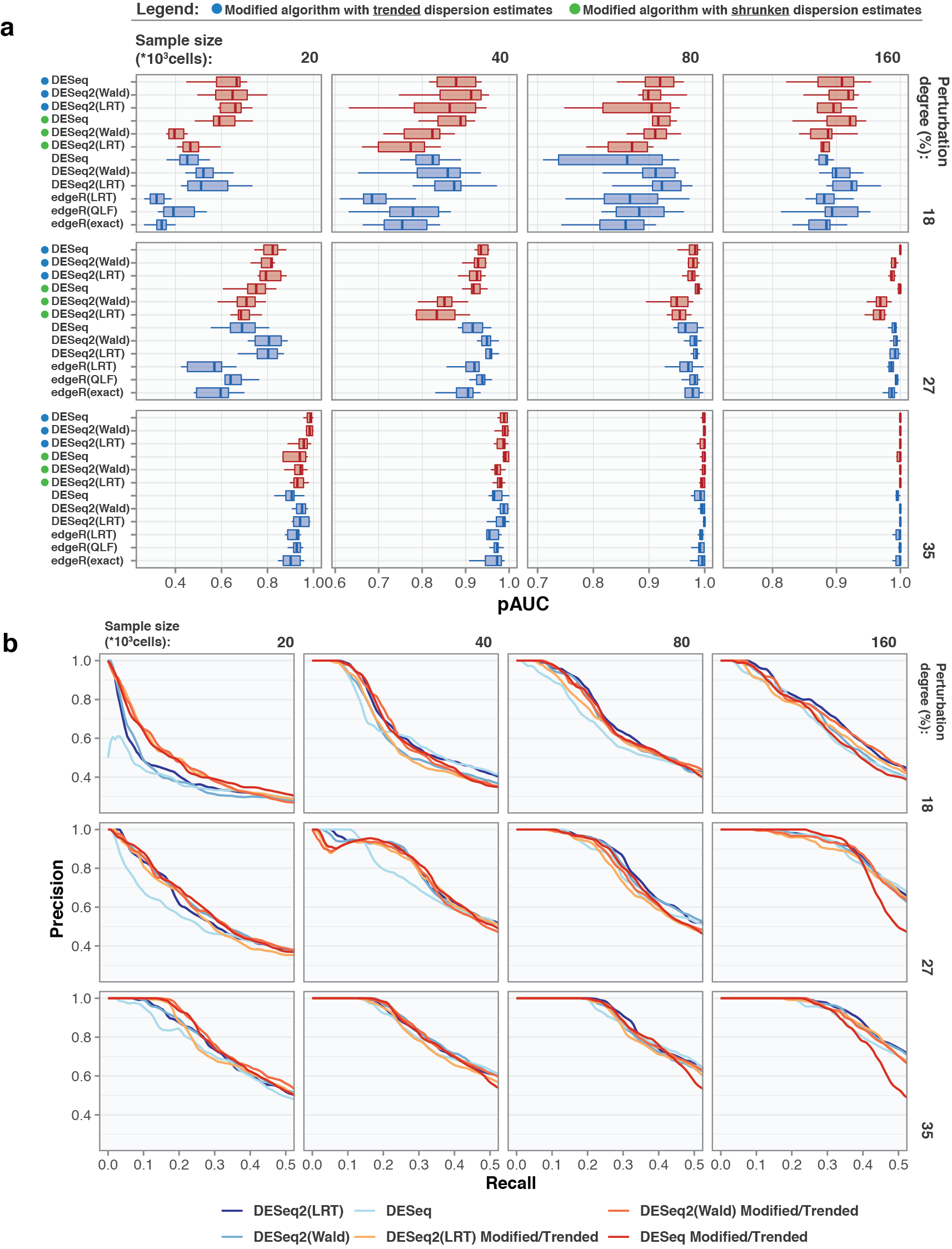

Supplementary Fig. 6. Comparison of the DRB scoring performance using precision-recall metrics.

Circles left to the algorithms’ names indicate the modified algorithms. **a**, Standardized partial area under the precision-recall curve (pAUC) calculated within intervals of [0,1] and [0,0.25] for precision and recall metrics, respectively. Shown are the pAUCs for perturbed subsamples of indicted size (top) and perturbation degree (right), with enriched to depleted barcodes ratio of 0.15. **b**, Precision-recall curves for the indicated sample sizes (top) and perturbation degrees (right), with enriched to depleted barcodes ratio of 0.15. For clarity, the modified algorithms with shrunken dispersion estimates are not shown.

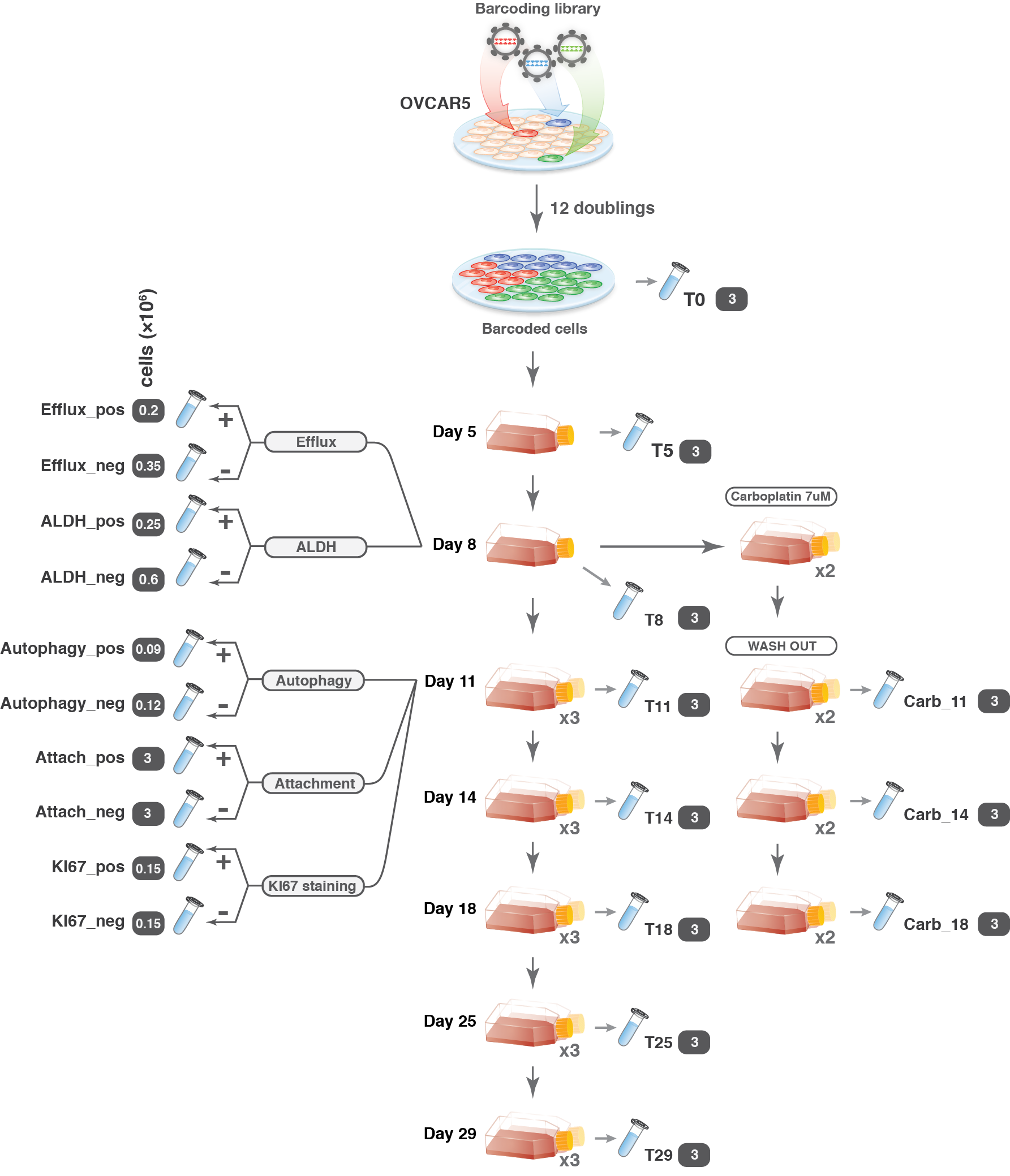

Supplementary Fig. 7. A schematic representation of OVCAR5 single-lineage phenotypic profiling experiment.

In the proliferation assay, 3 million cells were plated in each passage, and at day 8 population was split into 4 replicas. The tubes indicate that the sample(s) was collected for sequencing. Sample name as used in the read counts table (Supplementary Data 4) and number of cells collected (in millions; grey box) are marked near the tube images.

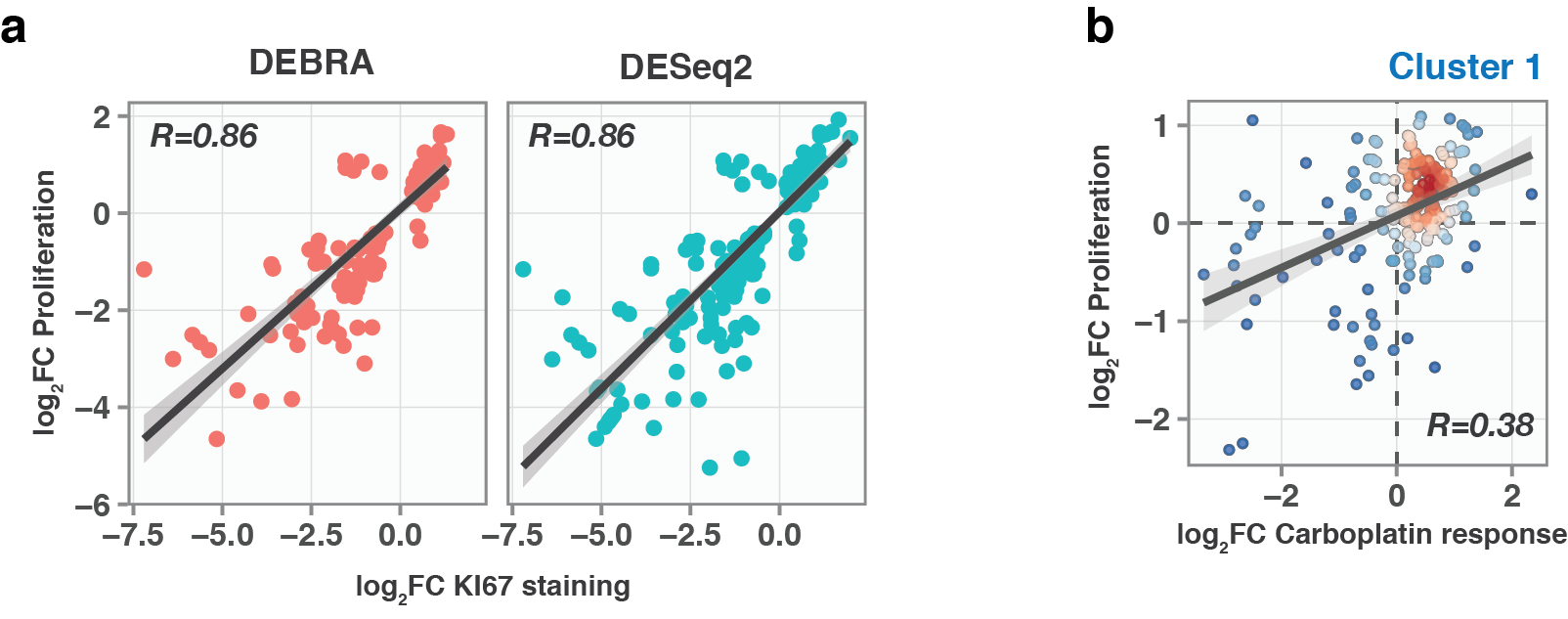

Supplementary Fig. 8. Barcode fraction fold-changes for OVCAR5 single-lineage phenotypes

**a**, Scatter plot of barcode fraction fold changes after 7 days growth and in KI67^HIGH^ cells compared to bulk analyzed with DEBRA and DESeq2 at FDR<0.25.

**b**, Scatter plot of barcode fraction fold changes in OVCAR5 cells in response to carboplatin treatment and after 7 days growth assay for cluster 1 (Bulk; see Fig. 5A).

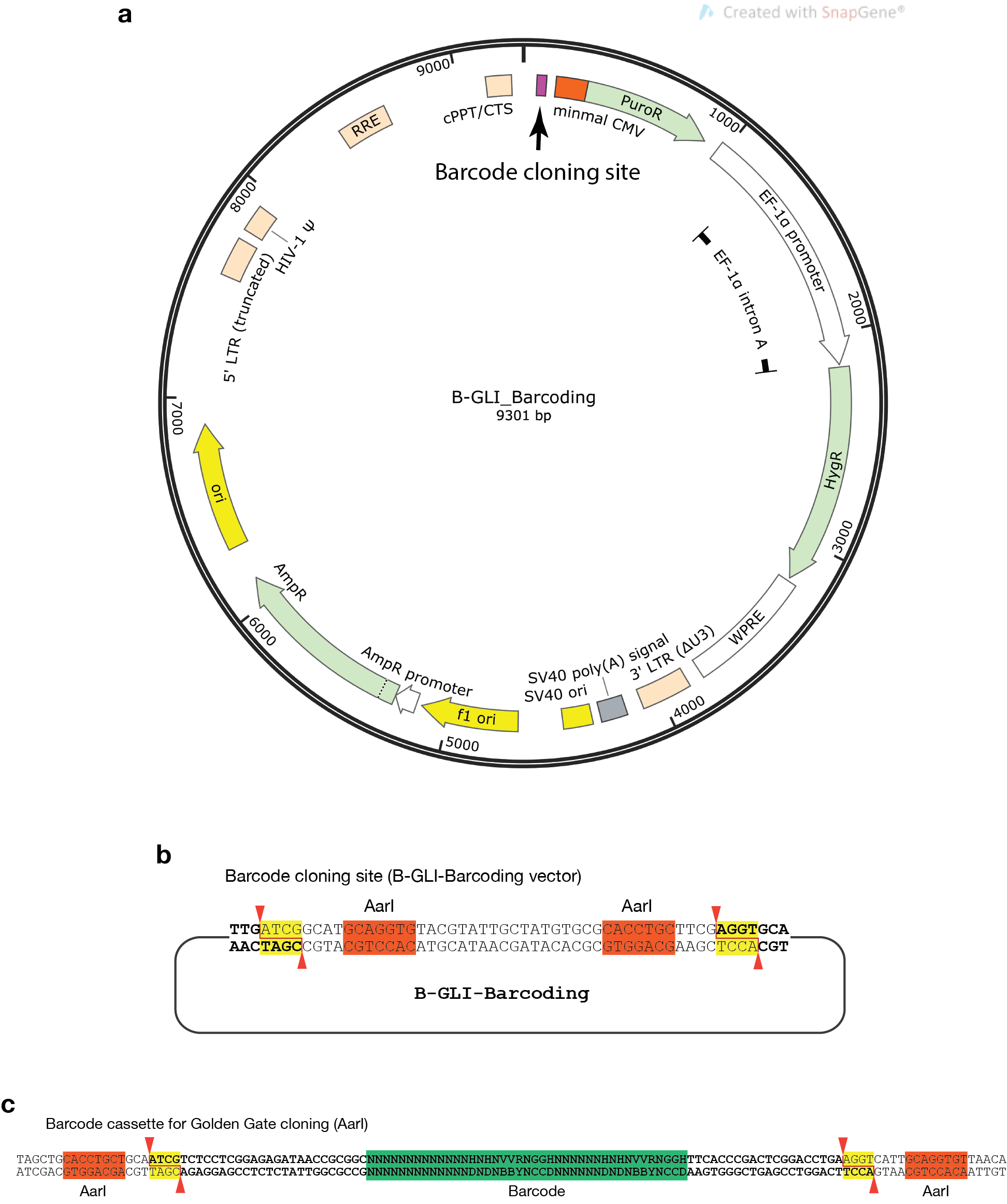

Supplementary Fig. 9. Lentiviral barcoding plasmid and cloning strategy outline

**a**, B-GLI-Barcoding plasmid map; image generated using SnapGene software (from GSL Biotech; available at snapgene.com)

**b**, Cloning site of B-GLI-Barcoding plasmid. AarI cut sites are marked in yellow.

**c**, Amplified barcode cassette as used for barcode cloning into B-GLI-Barcoding vector.

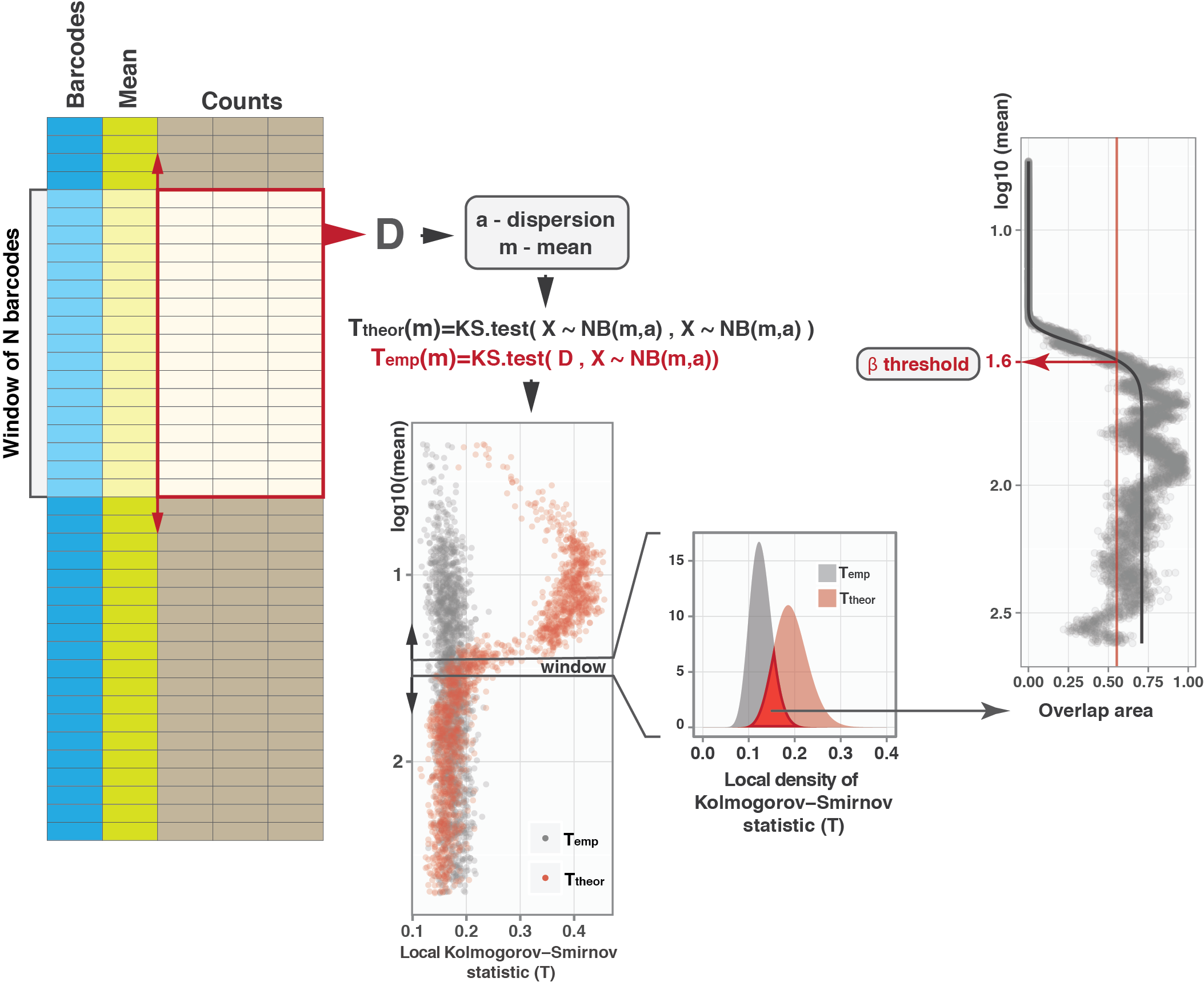

Supplementary Figure 10. The steps of the **β** threshold estimation algorithm.

The algorithm samples read count data ordered by the mean count value to obtain local estimates of mean and dispersion parameters of the negative binomial (NB) model. For each sampling window, theoretical and empirical Kolmogorov-Smirnov (KS) statistic values are calculated. Theoretical KS statistics is estimated on two sets of random negative binomial variables, whilst the empirical KS statistic is obtained by testing sampled data vs random negative binomial variables simulated using the previously estimated NB distribution parameters. After local estimation of these KS statistic values, the algorithm performs a local fitting of the empirical and theoretical KS statistic with a Gamma-distribution, and the overlap of local densities is fitted as a function of the mean read count using a 4-parameter sigmoid function. The **β** threshold is the value of the mean at which the fitted overlap takes the value of 0.8 of maximum.

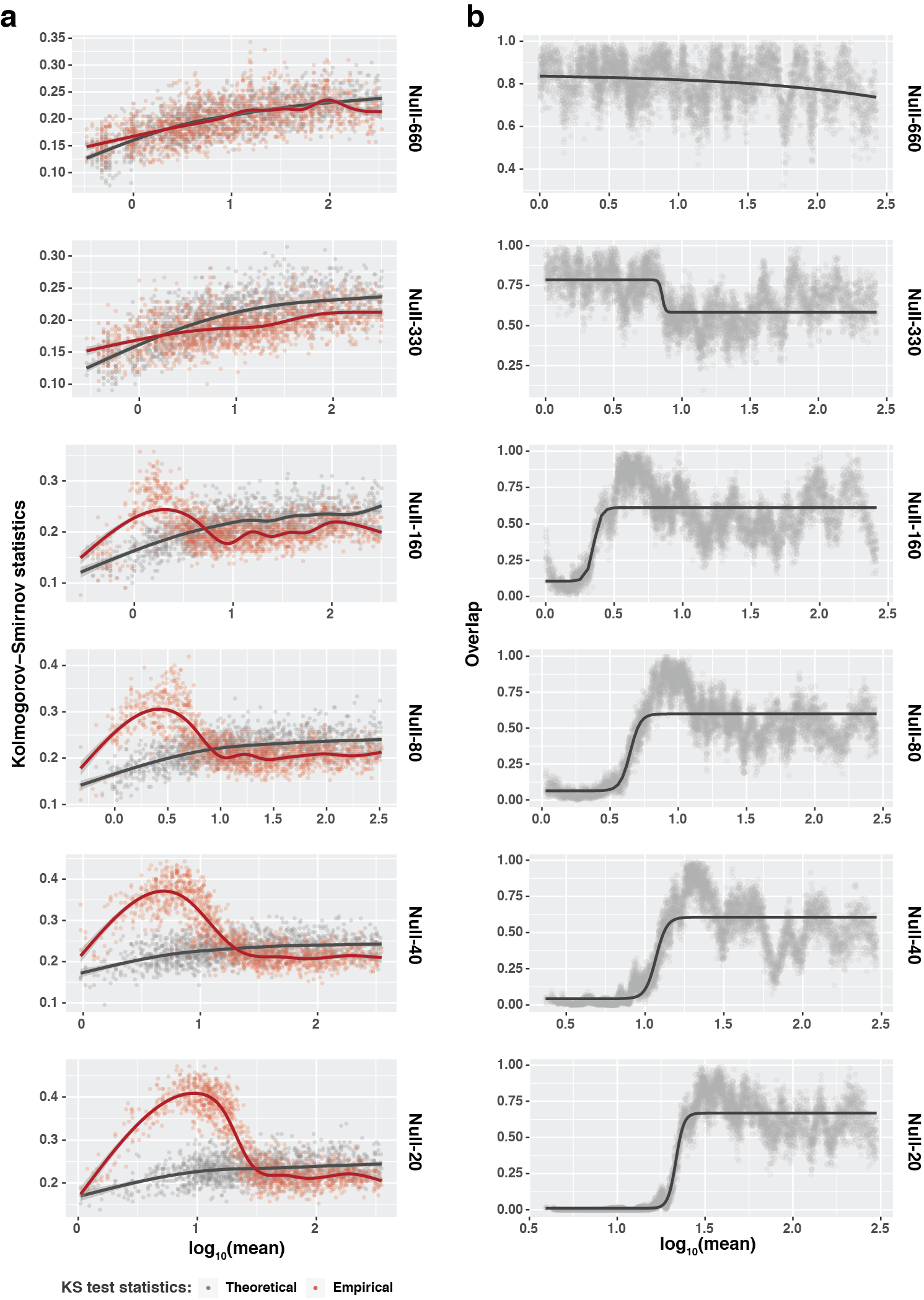

Supplementary Fig. 11. **β** threshold estimation algorithm applied to the null subsamples.

**a**, Empirical and theoretical (see Methods) local two-sample Kolmogorov-Smirnov test statistics for the null subsamples of different sizes. **b**, Estimation of the local overlap between Gamma-distributed empirical and theoretical Kolmogorov-Smirnov test statistics fitted with four parameters sigmoid function.

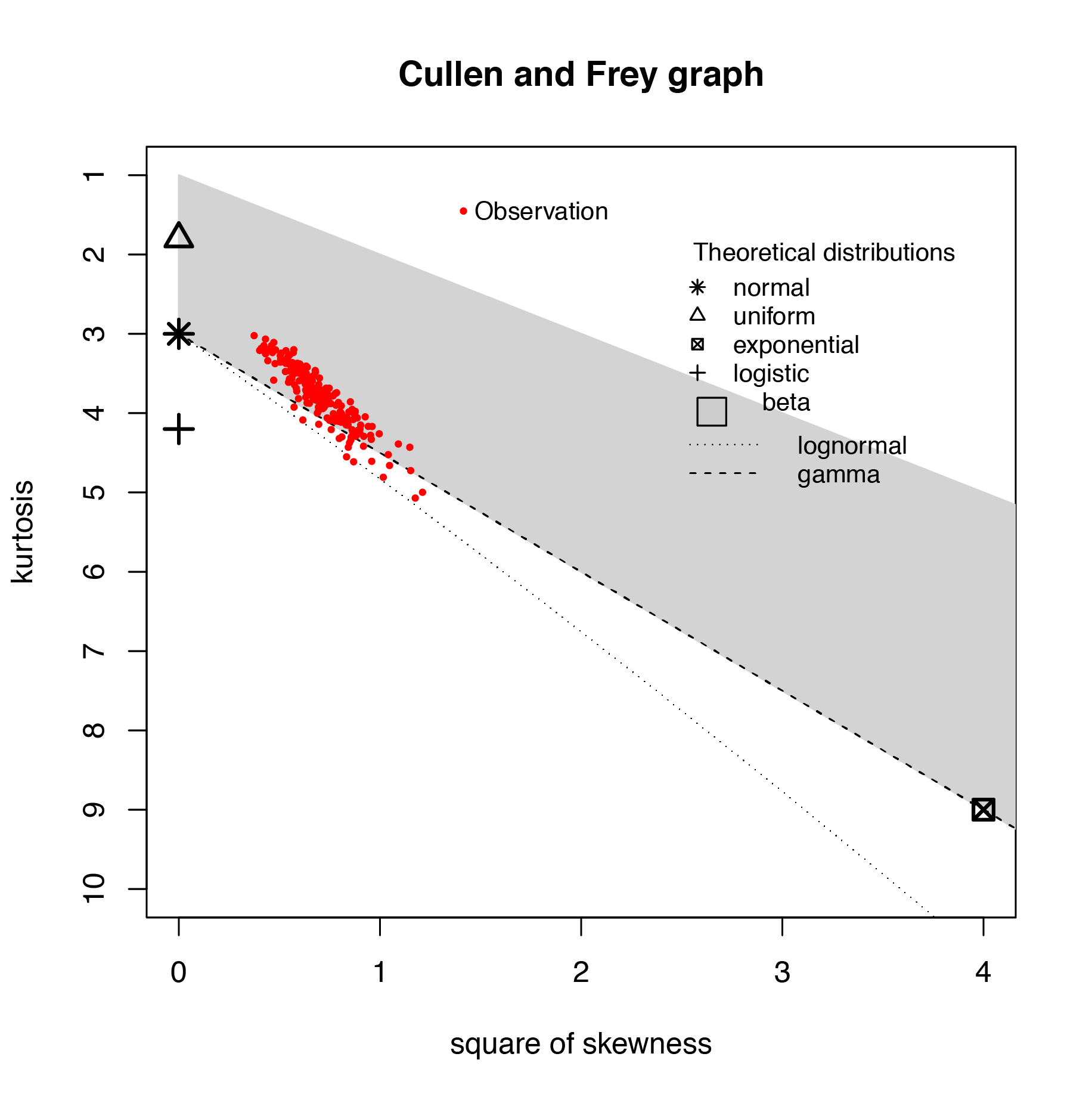

Supplementary Fig 12. Cullen and Frey graph for testing selected distributions of the Kolmogorov-Smirnov (KS) test statistics. Each red dot represents distribution of KS test statistics between two NB random variables with distinct mean and dispersion parameters obtained from 1000 resamples. The mean and dispersion values used for generating the random variables were derived from mean-variance modelling of null-80 samples using DESeq2.
